# Discovery of GS-5245 (Obeldesivir), an Oral Prodrug of Nucleoside GS-441524 that Exhibits Antiviral Efficacy in SARS-CoV-2 Infected African Green Monkeys

**DOI:** 10.1101/2023.04.28.538473

**Authors:** Richard L. Mackman, Rao Kalla, Darius Babusis, Jared Pitts, Kimberly T. Barrett, Kwon Chun, Venice Du Pont, Lauren Rodriguez, Jasmine Moshiri, Yili Xu, Michael Lee, Gary Lee, Blake Bleier, Anh-Quan Nguyen, B. Michael O’Keefe, Andrea Ambrosi, Meredith Cook, Joy Yu, Elodie Dempah, Elaine Bunyan, Nicholas C. Riola, Xianghan Lu, Renmeng Liu, Ashley Davie, Tien-Ying Hsiang, Michael Gale, Anita Niedziela-Majka, Joy Y. Feng, Charlotte Hedskog, John P. Bilello, Raju Subramanian, Tomas Cihlar

## Abstract

Remdesivir **1** is an amidate prodrug that releases the monophosphate of nucleoside GS-441524 (**2**) into lung cells thereby forming the bioactive triphosphate **2-NTP**. **2-NTP**, an analog of ATP, inhibits the SARS-CoV-2 RNA-dependent RNA polymerase replication and transcription of viral RNA. Strong clinical results for **1** have prompted interest in oral approaches to generate **2-NTP**. Here we describe the discovery of a 5’-isobutyryl ester prodrug of **2 (**GS-5245, Obeldesivir, **3**) that has low cellular cytotoxicity and three to seven-fold improved oral delivery of **2** in monkeys. Prodrug **3** is cleaved pre-systemically to provide high systemic exposures of **2** that overcome its less efficient metabolism to **2-NTP** leading to strong SARS-CoV-2 antiviral efficacy in an African green monkey infection model. Exposure-based SARS-CoV-2 efficacy relationships resulted in an estimated clinical dose of 350-400 mg twice-daily. Importantly, all SARS-CoV-2 variants remain susceptible to **2** which supports development of **3** as a promising COVID-19 treatment.

## Introduction

Wide-spread SARS-CoV-2 infection has resulted in the current COVID-19 pandemic, the largest global pandemic since the Spanish Flu, with more than 750 million reported cases and almost 7 million deaths.^1^ Multiple vaccines have been developed to combat the pandemic but due to the emergence of SARS-CoV-2 variants and continued transmission of the virus, the risk of hospitalization and severe disease remains a concern. Aside from vaccines, several treatment options have emerged including small molecule antivirals and antibodies that target different SARS-CoV-2 proteins to block viral replication and spread.^2–4^ Small molecule antivirals include the nucleotide analog remdesivir (**1**, Veklury®, Figure 1), and the nucleoside analog molnupiravir, that both target the viral RNA-dependent RNA polymerase (RdRp) by different mechanisms.^5–7^

**Figure 1.**
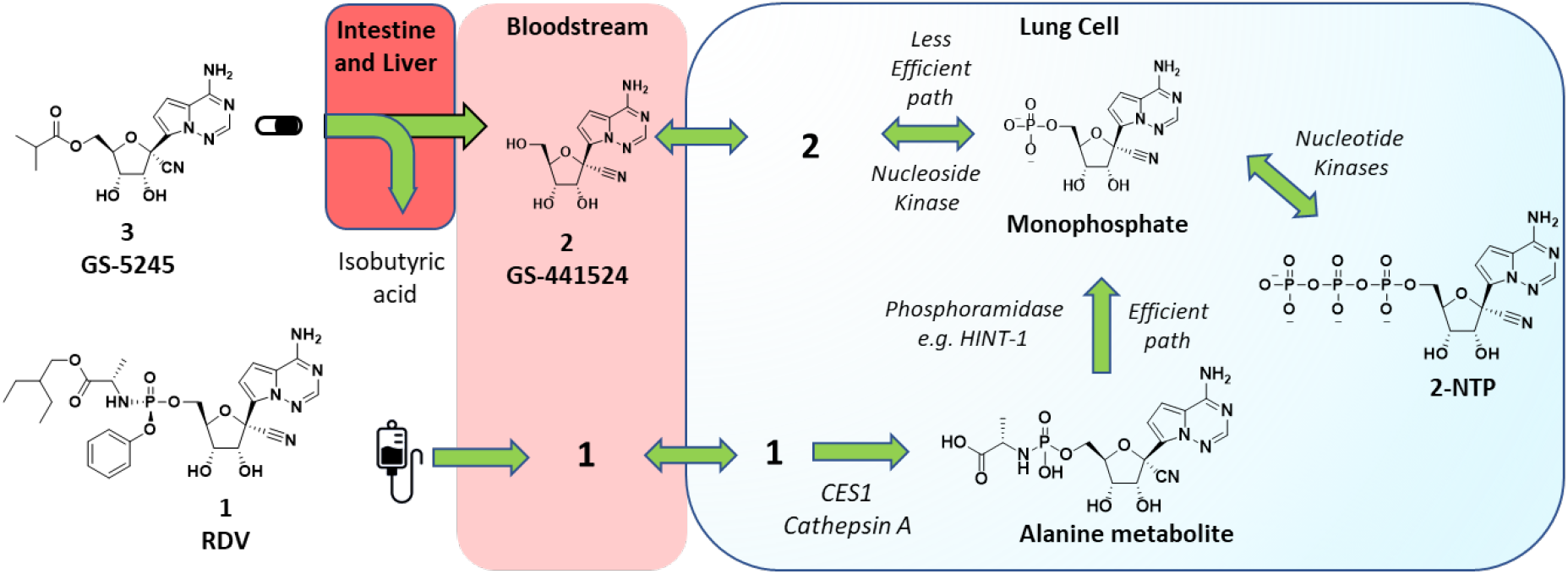
Metabolic activation pathways of prodrug **3** and phosphoramidate **1** to the common active **2-NTP** metabolite in lung. Ester prodrug **3** administered orally is metabolized pre-systemically in the intestine and liver to parent nucleoside **2** which then distributes into cells, including lung cells, where it is metabolized by a nucleoside kinase to the monophosphate. The monophosphate is then metabolized to the active metabolite, **2-NTP**, by the action of nucleotide kinases. These steps are reversible and allow the phosphorylated metabolites generated inside all cells to also be broken down to parent nucleoside **2** and released back into systemic circulation. Phosphoramidate **1**, following IV administration, rapidly distributes into many cell types, including lung cells, where it is metabolized irreversibly by the action of hydrolases (CES1, Cathepsin A) to the alanine metabolite and then to the same monophosphate by the action of phosphoramidases e.g. HINT-1. The metabolism of **1** to the monophosphate inside cells is efficient and is the major pathway for generation of **2-NTP** following administration of **1**.

Nirmatrelvir is an inhibitor of the SARS-CoV-2 3CL main protease and is administered in combination with the P450 3A4 inhibitor ritonavir as a pharmacokinetic enhancer (combination product marketed as Paxlovid®).^8^ In the outpatient setting, 5-day oral Paxlovid and 3-day intravenous (IV) **1** treatment have both demonstrated ≥87% effectiveness in reducing the hospitalization of high-risk SARS-CoV-2 infected patients.^9, 10^ Despite the promising efficacy of these treatment options, limitations in their broad application exist. Nucleotide prodrug **1** requires IV administration, molnupiravir has safety concerns due to its mutagenic mechanism of action, and oral Paxlovid is contraindicated in some patients due to drug-drug interactions.^7, 11^ These drawbacks indicate continued exploration to discover novel oral treatment options for COVID-19 is imperative.

Nucleotide **1** is the first approved treatment for COVID-19 and is a chiral phosphoramidate prodrug bearing the 2-ethylbutyl-*L*-alanine and phenol pro-moieties.^12^ The pro-moieties are intracellularly cleaved to generate the monophosphate of the parent nucleoside **2** in different cells and tissues including lung cells (Figure 1). Initially, the parent nucleoside **2** was designed as a potential treatment for hepatitis C virus but was then found through broad screening to be more potent toward respiratory syncytial virus (RSV).^12–15^ Phosphate prodrug exploration aimed at improving the formation of the active metabolite **2-NTP** in lung cells for RSV then resulted in the initial discovery of **1**.^12^ Following IV administration the phosphoramidate prodrug **1** rapidly distributes into lung cells, in addition to other tissues, where it is efficiently metabolized by enzymes into the monophosphate nucleotide, effectively bypassing a rate-limiting first phosphorylation step of **2** to the same monophosphate intermediate (Figure 1). The monophosphate is then further metabolized to the active **2-NTP** that inhibits the viral replication process of multiple viral polymerases including HCV, RSV, Dengue, Ebola, Nipah, SARS-CoV, MERS and more recently, SARS-CoV-2.^5–6, 16–17^ Prior to the pandemic, efficacy studies in animal models of RSV, and SARS-CoV and MERS coronaviruses, supported the potential utility of **1** as a treatment for these respiratory viruses.^12, 18–20^ In the first clinical studies in patients hospitalized with COVID-19, IV administration of **1** was shown to shorten the hospitalization duration by 5 days (ACTT-1 trial) and improve survival of hospitalized patients with COVID-19.^21–23^ A more recent clinical study in high-risk non-hospitalized patients (PINETREE trial) demonstrated an 87% reduction in hospitalizations following a short 3-day IV administration of 1.^9^ This result indicates that delivery of the active **2-NTP** metabolite within the lung during an early stage of SARS-CoV-2 infection can provide a meaningful clinical benefit for patients.

The clinical evidence for **1** as a COVID-19 treatment has prompted much interest in its potential for oral delivery. Unfortunately, the metabolic process that occurs in lung cells to generate the **2-NTP** metabolite also occurs in other tissues, including the liver, which effectively hydrolyzes **1** following oral delivery. In non-human primates (NHPs), <1% oral bioavailability of the intact prodrug **1** was observed suggesting that orally delivered **1** would be ineffective at providing sufficient intact prodrug in systemic circulation that is comparable to IV administration in order to be efficacious.^12^ Therefore, shortly after the pandemic started, we initiated programs aimed at orally delivering effective concentrations **2-NTP** into lungs leading to the selection of the 5’-isobutyryl ester prodrug **3 (**GS-5245, Obeldesivir, ODV) as a clinical candidate. Here we disclose the chemical structure, synthesis, and in vitro biophysical evaluation of **3** together with its oral properties and efficacy in the SARS-CoV-2 infection model in African green monkeys (AGM). Importantly, the efficacy of **3** in the AGM model closely matched that of **1** which was dosed at clinically relevant exposures in the same AGM model, suggesting that oral **3** has the potential to match the efficacy of **1** in the clinic.^24^ In addition to this work, other groups have identified and characterized prodrugs of **2**, including a 5’-isobutyryl ester **3**.^25^ Efficacy was reported across several mouse models but no plasma exposure-efficacy analysis was reported to allow human doses to be estimated. Here we utilized exposure-efficacy data for **3** from the AGM model together with data from several independent SARS-CoV-2 models with an earlier prodrug **6** to develop an exposure-SARS-CoV-2 efficacy model. This model was then used to estimate a human daily dose of 3 that ranged from 350-400 mg twice-daily. The results presented here supported the rapid advancement of **3** into the clinic as a potential oral treatment for COVID-19.

## Results and Discussion

In order to generate the **2-NTP** metabolite from an orally delivered compound we considered other monophosphate prodrugs besides **1** that had improved in vitro liver stability in addition to the parent nucleoside **2**, which has a long human microsomal half-life of >8 h in vitro (Table 1).

**Table 1.**
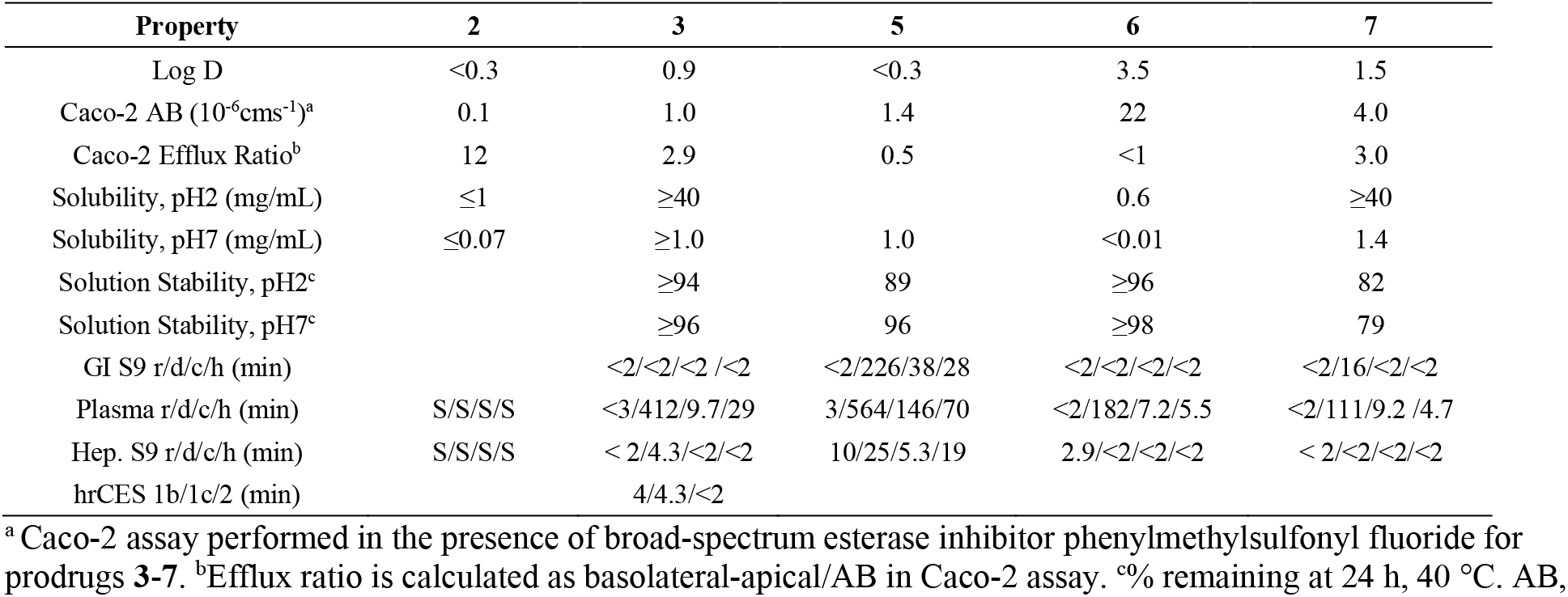
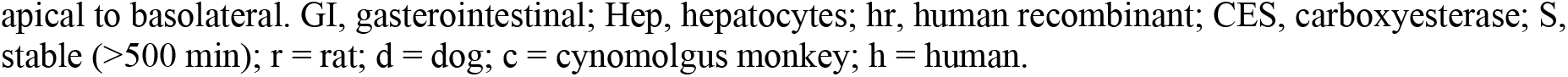
In vitro pharmacokinetic and physical properties of **2** and its prodrugs **3**, **5**-**7**.

Following IV administration in NHPs **2** demonstrated an in vivo half-life of 2.6-2.7 h which is significantly longer than the 0.4-0.8 h previously reported for amidate **1** in NHPs (Table 2).^12^

**Table 2.** In vivo plasma pharmacokinetic parameters of **2** and **3**^a^.

| Route /Cpd. | Model | Dose<sup>b</sup> | **2**<br>CL<br>(L/h/kg) | **2**<br>V<sub>ss</sub><br>(L/kg) | **2**<br>T<sub>1/2</sub><br>(h) | **2**<br>C<sub>max</sub><br>(μM) | **2**<br>AUC<sub>0-24</sub><br>(μM.h) | **3**<br>AUC<sub>0-24</sub><br>(μM.h) | Lung<br>[2-NTP]<br>(nmol/g) | **2**<br>F% | **2**<br>F%<br>(Solid) |
| --- | --- | --- | --- | --- | --- | --- | --- | --- | --- | --- | --- |
| **IV**<br>**2**<sup>c</sup> | Rat | 1 | 1.36 ± 0.02 | 1.66 ± 0.06 | 1.03 ± 0.02 | 1.78 ± 0.28 | 2.50 ± 0.04 |  |  |  |  |
|  | Dog | 1 | 0.18 ± 0.01 | 0.33 ± 0.06 | 1.82 ± 0.11 | 13.5 ± 2.9 | 18.7 ± 1.5 |  |  |  |  |
|  | Cyno | 20 | 0.56 ± 0.05 | 1.03 ± 0.09 | 2.68 ± 0.07 | 79.2 ± 6.7 | 123 ± 11.1 |  | 0.41 ± 0.09 |  |  |
|  | AGM | 20 | 0.49 ± 0.12 | 0.89 ± 0.26 | 2.63 ± 0.19 | 87.3 ± 37.4 | 147 ± 36 |  | 0.23 ± 0.13 |  |  |
| **Oral**<br>**2**<sup>c</sup> | Rat<sup>d</sup> | 5 |  |  | 1.87 ± 0.59 | 1.11 ± 0.30 | 3.97 ± 1.19 |  |  | 32 ± 10 |  |
|  | Dog | 5 |  |  | 4.18 ± 0.64 | 27.3 ± 3.5 | 83.4 ± 14.4 |  |  | 89 ± 15 |  |
|  | Cyno | 5 |  |  | 3.31 ± 0.65 | 0.54 ± 0.28 | 1.67 ± 0.76 |  |  | 5.4 ± 2.5 |  |
|  | AGM | 5 |  |  | 6.65 ± 1.82 | 0.36 ± 0.05 | 3.15 ± 0.94 |  |  | 8.6 ± 2.5 |  |
| **Oral**<br>**3**<sup>c</sup> | Rat | 6 |  |  | 2.02 ± 0.33 | 2.10 ± 0.49 | 7.57 ± 0.53 | BLQ |  | 63 ± 4 | 90 ± 15 |
|  | Dog | 14.4 |  |  | 3.62 ± 0.23 | 57.8 ± 11.3 | 202 ± 44 | 0.40 ± 0.04 |  | 93 ± 20 | 67 ± 12 |
|  | Cyno | 11.7 |  |  | 2.62 ± 0.55 | 7.57 ± 6.07 | 21.6 ± 11.0 | 0.01 ± 0.01 |  | 37 ± 19 | 18 ± 3 |
|  | AGM | 14.4 |  |  | 3.19 ± 0.19 | 6.81 ± 3.08 | 26.6 ± 12.1 | 0.02 ± 0.02 |  | 31 ± 14 | - |
|  | AGM | 60 |  |  | 2.90 ± 0.04 | 22.5 ± 1.3 | 129 ± 2 | 0.03 ± 0.03 | 0.31 ± 0.09 | 36 ± 1 |  |
|  | AGM | 86.5 |  |  | 2.98 ± 0.22 | 19.6 ± 9.8 | 136 ± 67 | 0.68 ± 0.75 | - | 27 ± 13 |  |

<sup>a</sup>Doses and parameters shown are for the solution formulation. All studies are n=3 animals unless noted. <sup>b</sup>mg/kg.
<sup>c</sup>Formulations described in the experimental methods. <sup>d</sup>n=6 animals. Cyno, cynomolgus monkey; AGM, African green monkey.

Allometric scaling of the IV data resulted in a projected human half-life for **2** of ∼4 h which supported a convenient twice-daily regimen for sustained plasma exposures of **2**. Despite the promising liver stability of **2**, some significant challenges hindering its oral potential were identified. The first challenge was its inefficient intracellular metabolism from **2** to its monophosphate and then to **2-NTP** as described in Figure 1.^12^ This less efficient metabolism has also been reported by independent groups and manifests as weaker antiviral activity across different lung cells in vitro compared to **1**.^12, 24, 26^ The reduced **2-NTP** formation in vitro was also observed in vivo through analyzing lung tissue samples harvested at 24 h from NHPs and ferrets dosed IV with either **2** or **1**.^12, 24, 27^ These results suggest that high plasma exposures of **2** are likely necessary to drive sufficient lung **2-NTP** formation in vivo to achieve comparable efficacy to **1**. The second challenge was the poor physicochemical properties of **2** including a low apical to basolateral permeability in Caco-2 cells, a 12-fold efflux ratio, and low pH 7 solubility (Table 1). These physicochemical properties result in a low and variable oral bioavailability across preclinical species ranging from <8.6% in two species of NHPs to 89% in dog (Table 2). The high oral bioavailability in dogs was not considered to be a reliable predictor of human oral potential given the low F% in NHPs and the ‘leaky intestine’ phenomena in dogs as a result of their larger and more abundant paracellular junctions compared to human.^28^ Indeed, the 4’-azido cytidine ribonucleoside analog R1479 had a similar preclinical profile in NHPs and dogs to that of **2**, but in humans only an estimated 6-18% oral bioavailability was observed consistent with the NHP data.^29^ Our strategy therefore focused on addressing the oral limitations of **2** with prodrugs that could effectively deliver high concentrations of **2** to overcome its metabolic deficiencies and drive efficacy. Finally, nucleoside analogs such as **2** are challenging and expensive to manufacture due to their structural complexity. The prospect that a prodrug could significantly reduce the proportion of the administered dose that is eliminated without absorption was highly appealing as a strategy to reduce the likely dose required.

Scheme 1 describes the synthesis for several ester-based prodrugs including 5’-mono esters **3** and **5**, and the tri-esters **6** and **7.**

**Scheme 1.**
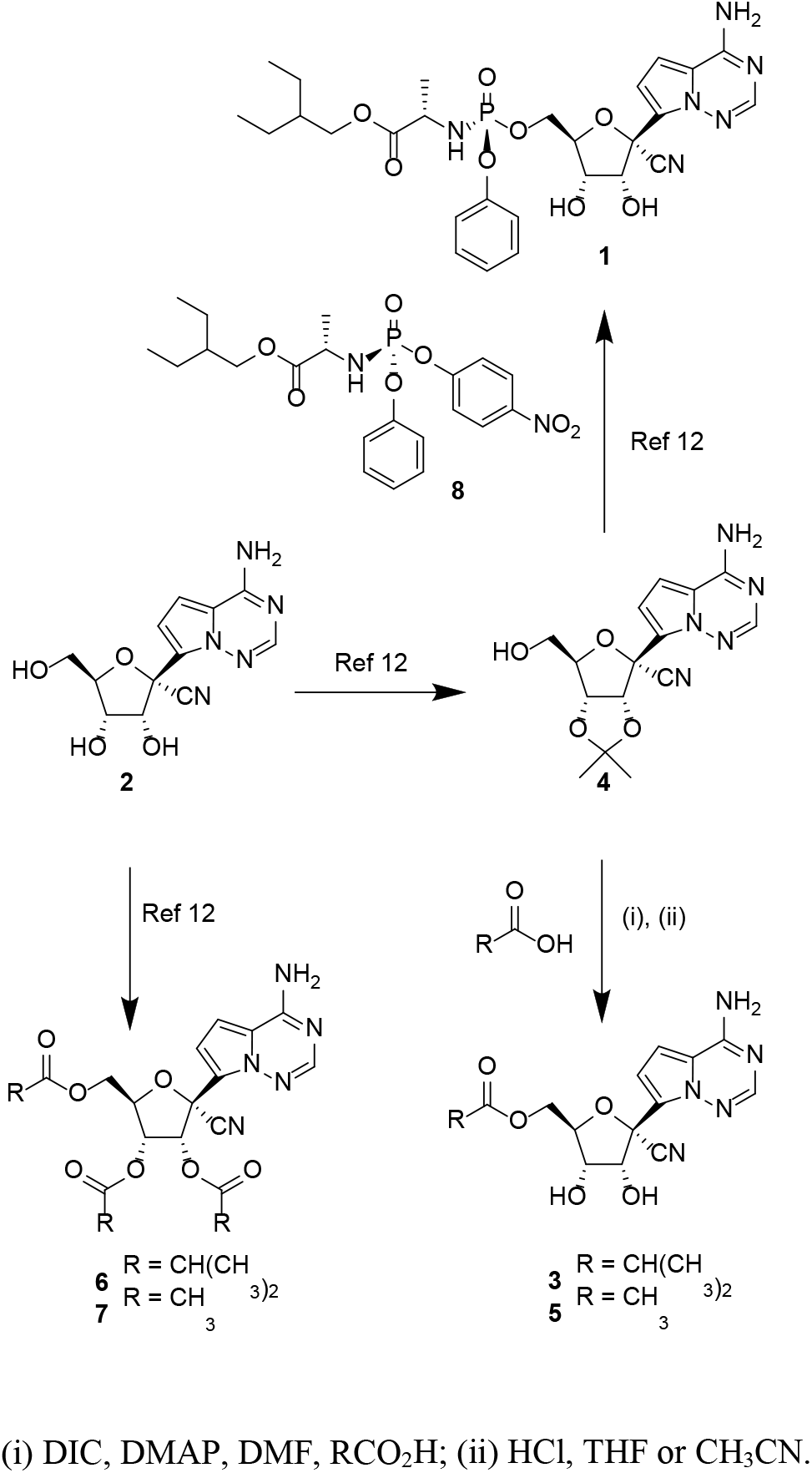
Synthesis of 5’-ester prodrugs.

The latter two prodrugs were prepared prior to the pandemic as potential oral treatments for RSV and their synthesis has been previously reported.^12^ The 5’-mono ester prodrugs were readily synthesized from the 2’,3’-acetonide protected nucleoside **4**, a known intermediate within our remdesivir synthetic process.^30^ Straightforward coupling of the intermediate **4** with alkyl carboxylic acids using carbodiimide was followed by acid-mediated cleavage of the 2’,3’-protecting group to yield the monoesters **3** and **5,** respectively. Of note, amidate **1** is also synthesized from the same intermediate **4** through a similar two-step process, but instead utilizes a synthetically complex, chiral 5’-mono-phosphoramidate prodrug reagent **8**.^30^ The reduced synthetic complexity of the ester prodrugs is a favorable feature that may lead to downstream benefits in scalability relative to amidate **1**. The ability to rapidly scale and deliver more cost-effective therapies for COVID-19 is important for broad accessibility. However, despite the reduced complexity of **3**, the per mole dose required for efficacy is likely to be significantly higher than the dose of **1** which will limits the impact of its reduced complexity.

The permeability of the ester prodrugs was compared to parent **2** using a Caco-2 cell assay in the presence of bis(nitrophenol) phosphate, a broad-spectrum carboxyesterase inhibitor to suppress rapid esterase-mediated cleavage of the prodrugs. Mono esters **3** and **5** both demonstrated ∼10-fold improved forward permeability over parent **2** and reduced the Pgp-mediated efflux ratio to <3-fold (Table 1). In contrast the tri-ester prodrug **6** increased the log D above 3 and significantly improved passive permeability with no evidence of efflux. As expected, the permeability properties of the esters trended in line with their log D, with the tri-esters **6** and **7** demonstrating the highest log D values and more favorable permeability properties over the mono esters.

The thermodynamic solubility of the esters in crystalline or amorphous solid forms was assessed at pH2 and pH7. The triester **6,** with the highest log D, demonstrated the lowest solubility and was less soluble than **2**. In contrast, the less lipophilic iso-butyryl ester **3** and acetate esters **5** and **7** all demonstrated >10-fold improved pH7 solubility over **2**, and >∼100-fold over **6**. Overall, the combined solubility and permeability of **3**, **5** and **7** was considered more favorable than **6** especially when formulating solid dosage forms where dissolution properties are more critical. Chemical pH-dependent stability was also assessed to determine the relative potential for the prodrugs to remain intact in the stomach and intestine during oral delivery. Ester **3** demonstrated excellent pH2 and pH7 stability while the acetate esters **5** and **7** were both less stable, suggesting **3** was the optimal compound based on overall stability, permeability and solubility.

Crystallization studies on **3** resulted in the identification of multiple crystalline forms, including stable salt-free Form III (see supporting information). The X-ray crystal structure of Form III confirmed the overall chemical structure of **3**. In contrast, solid crystalline forms of the tri-ester **6** were challenging to identify, and an extensive salt screening effort was required to identify the crystalline mono-pyruvate salt. In this example, the pyruvate salt combined with the tri-isobutyryl esters resulted in a significant 1.6-fold increase in molecular weight compared to **3** which was considered a disadvantage with respect to future pill mass. Hydrobromide salt forms of **6** have now been reported in addition to the pyruvate form.^31^ Taken together, the in-vitro assessments indicated that the overall physicochemical properties for the mono 5’-iso-butyryl ester **3** including permeability, solubility, pH-dependent stability, and crystalline form isolation, favored its selection as the candidate compound.

The cell-based SARS-CoV-2 antiviral activity of **3** and other ester prodrugs was evaluated in A549-hACE2 and NHBE cells but not considered part of our candidate selection process. This was because the intact prodrug was not expected to be present in vivo based on our earlier experience with ester prodrugs on **2** for RSV (see later discussion). The antiviral assays were however useful to dissect the factors governing the metabolism of **2** to **2-NTP** by intracellular kinases compared to the metabolism of **1** to **2-NTP** which occurs via a different, metabolic pathway (Figure 1). As shown in Table 3, **3** inhibits SARS-CoV-2 with an EC_50_ value of 1.90 ± 0.61 µM in the A549-hACE2 cell line and has comparable potency to **2**.

**Table 3.** SARS-CoV-2 Antiviral activity and cytotoxicity profiling of **3**.

| Antiviral Activity EC<sub>50</sub> (μM)<sup>a</sup> |  | 1 | 2 | 3 | Puromycin |
| --- | --- | --- | --- | --- | --- |
| A549-hACE2<sup>b</sup> | Lung carcinoma | 0.06 ± 0.03 | 2.49 ± 0.62 | 1.90 ± 0.61 |  |
| NHBE<sup>c</sup> | Primary | 0.04 ± 0.01 | 3.25 ± 0.8 | 0.43 ± 0.09 |  |
| Cytotoxicity CC<sub>50</sub> (μM)<sup>a</sup> |  |  |  |  |  |
| A549-hACE2 | Lung carcinoma | 22.9 ± 6.9 | >50 | >50 |  |
| Huh7 | Hepatoma |  | >89 | >44 | 0.43 ± 0.08 |
| HepG2 | Hepatoblastoma (galactose) | 11.1 ± 1.2<sup>d</sup> | >100<sup>d</sup> | 71 ± 7.5 | 0.78 ± 0.22 |
| PC-3 | prostate cancer (galactose) | 1.4 ± 0.1<sup>d</sup> | >100<sup>d</sup> | >100 | 0.47 ± 0.05 |
| MT-4 | lymphoblast T-cell | 1.7 ± 0.4<sup>d</sup> | 69 ± 26<sup>d</sup> | >50 | 0.12 ± 0.03 |
| MRC5 | lung fibroblast |  | >89 | >44 | 0.32 ± 0.03 |
| NHBE | primary | >50 | >50 | >50 | 0.60 ± 0.2 |
| Quiescent PBMCs | primary | >20<sup>d</sup> | >100<sup>d</sup> | >100 | 5.05 ± 0.01 |
| Stimulated PBMCs | primary | 14.8 ± 5.8<sup>d</sup> | >100<sup>d</sup> | >100 | 1.10 ± 0.01 |
| PHH | primary | 2.5 ± 0.6<sup>d</sup> | >100<sup>d</sup> | >100 | 1.1 ± 0.4 |

<sup>a</sup>Values are the mean ± standard deviation (SD) of the results from ≥3 independent experiments. <sup>b</sup>2-day SARS-CoV-2-Nluc antiviral assay. <sup>c</sup>1-day SARS-CoV-2-Fluc antiviral assay. <sup>d</sup>Reference 32.

This reflects the facile ability of **3** to break down to **2** in the cell culture experiment leading to the same potency. However, in NHBE cultures **3**, rather unexpectedly, demonstrated higher potency than **2** with an EC_50_ value of 0.43 ± 0.09 µM suggesting that **3** may be facilitating more rapid delivery of **2** into the NHBE cells because of its improved permeability properties. Amidate prodrug **1** exhibited significantly greater potency than either **2** or **3** against SARS-CoV-2 across both cell lines. In both A549-hACE2 and NHBE cell cultures, neither **2** or **3** were found to be cytotoxic at concentrations up to 50 μM and amidate **1** demonstrated at least 300-fold selectivity (CC_50_/EC_50_) (Table 3). To correlate antiviral potency with intracellular concentrations of **2-NTP** both cell lines were incubated continuously with **2** or **3** at 10 µM, or **1** at 1 µM and the **2-NTP** concentrations were determined by LCMS at different timepoints over 48 h (Figure 2A and 2B).

**Figure 2.**
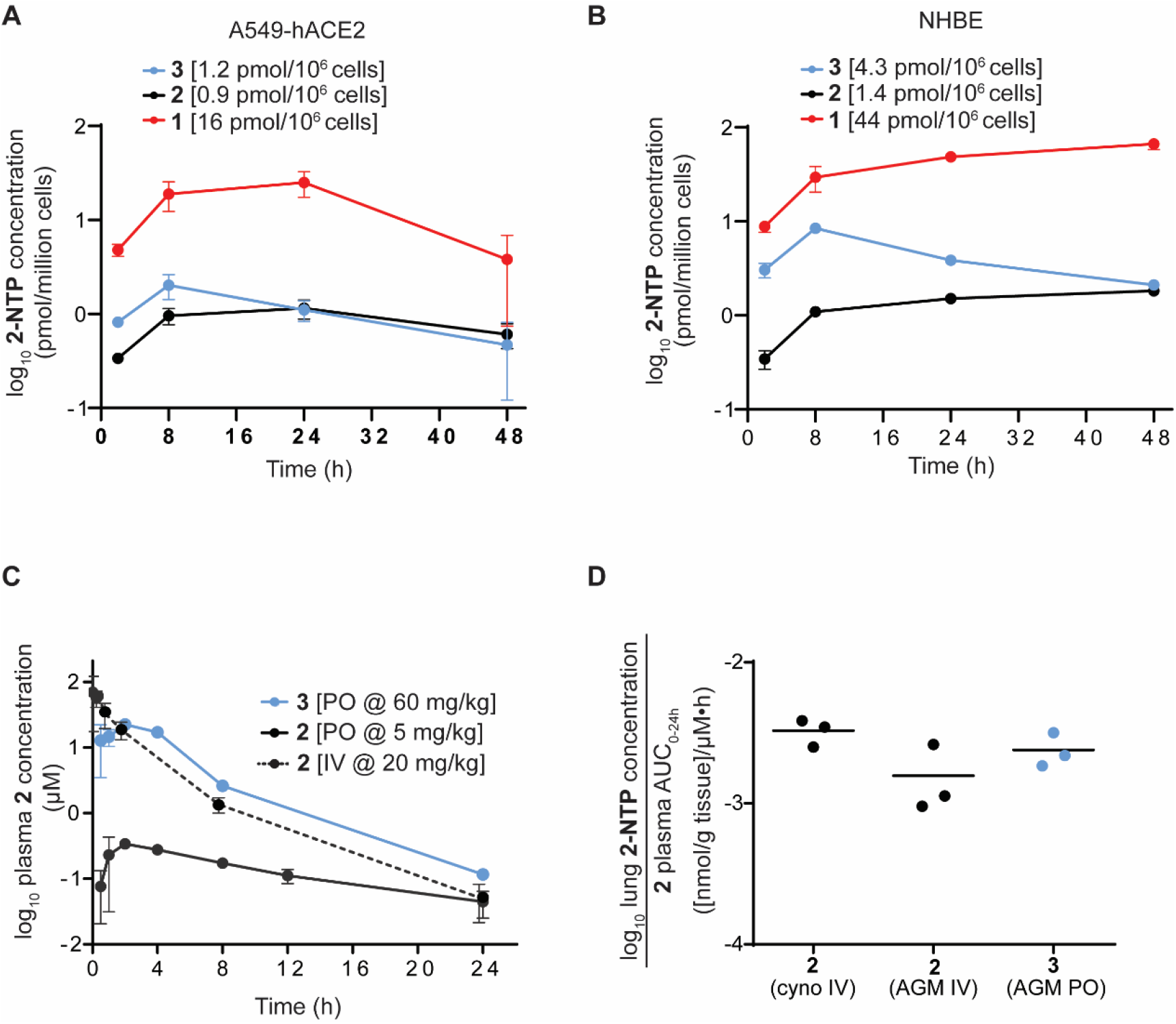
Cellular metabolism and in vivo pharmacokinetics of **3**. Panel **A**, Intracellular metabolism of **3** (blue), **2** (black), and **1** (red) to the active metabolite **2-NTP** in A549-hACE2 cells. Average **2-NTP** concentrations are indicated after dose-normalization to 1 µM. Panel **B**, Intracellular metabolism in NHBE cells. Average **2-NTP** concentrations are indicated after dose-normalization to 1 µM. Panel **C**, Plasma concentration-time profile of **2** following IV and oral administration and oral administration of **3** in AGM. Panel **D**, Lung **2-NTP** concentrations at 24 h normalized to exposure of **2** in plasma following IV administration of **2** in both cynomolgus monkey and AGM, and oral dosing of **3** in AGM.

In both human immortalized and primary human lung cells, **2** yielded similar average concentrations of **2-NTP** consistent with its comparable antiviral activity in the cell cultures. The **2-NTP** concentrations from **2,** when normalized to the same incubation concentration of **1,** were >17-fold lower reflecting the weaker antiviral activity of **2** relative to **1** and confirming the greater efficiency of the amidate prodrug for generating the common monophosphate intermediate, and ultimately **2-NTP** in these cell cultures. For prodrug **3** the average **2-NTP** concentrations were also consistent with the antiviral activity profile. Similar average **2-NTP** concentration were observed in the A549-hACE2 cells where antiviral activity was comparable, but approximately 3-fold elevated concentrations were noted in the NHBE cells from **3** especially at the early timepoints which drives the higher potency. In summary, the average **2-NTP** concentrations align very well with the antiviral data and confirm that **2** is rate-limited in its first intracellular metabolism step in lung cells to the monophosphate relative to the amidate prodrug **1**. Moreover, poor cell permeability is also implicated as a factor contributing to the lower efficiency for generating **2-NTP** from **2**.

Following oral dosing, cells in the intestine, liver, and circulating immune cells are expected to be exposed to varying concentrations of **3** over time in addition to high systemic concentrations of **2**. Prodrug **3** and the main metabolite **2** were therefore evaluated for toxicity across a panel of five human transformed cell lines and several primary human cell types, including quiescent and stimulated peripheral blood mononuclear cells (PBMCs), and primary human hepatocytes (PHH) with a 5-day treatment (Table 3). Low toxicity of **3** was observed with CC_50_ values >44 µM, consistent with parent **2** that also showed minimal toxicity in all cells tested. Prodrug **3** was also found to be non-toxic in PHH and PBMCs up to 100 µM and neither **3** nor **2** showed any inhibition of mitochondrial respiration and mitochondrial protein synthesis at the highest concentration tested (100 µM) (See supporting information).^32^ Similarly, a minimal effect was observed on the mitochondrial DNA synthesis when the cells were treated with 0.4-40 µM **3** or 1.0-100 µM **2**. Taken together these data suggest that **3** and its metabolite **2** have low risks for cellular and mitochondrial toxicity in relevant tissues that would be exposed to these compounds upon oral delivery.

Since the start of the pandemic, multiple new variants of SARS-CoV-2 have emerged. Sequence analysis suggests that the majority of amino acid substitutions observed in variants occur in the envelope glycoprotein (spike), while the SARS-CoV-2 RdRp has been highly conserved.^33^ Due to this high conservation, the active **2-NTP** metabolite of **1** and **2** has been shown to retain antiviral activity against emergent variants up to the Omicron variant (B.1.1.529/BA.1).^34^ Here we assessed the potency of **3** and against a broad panel of new variants of concern (VOC) including Omicron and its subvariants, and the Delta variant.^35^ Activity was assessed in an antiviral ELISA assay performed in the A549-hACE2-TMPRSS2 cell line and the results are reported in Table 4 along with **1** and **2** as fold change relative to the ancestral WA1 isolate (Lineage A).

**Table 4.**
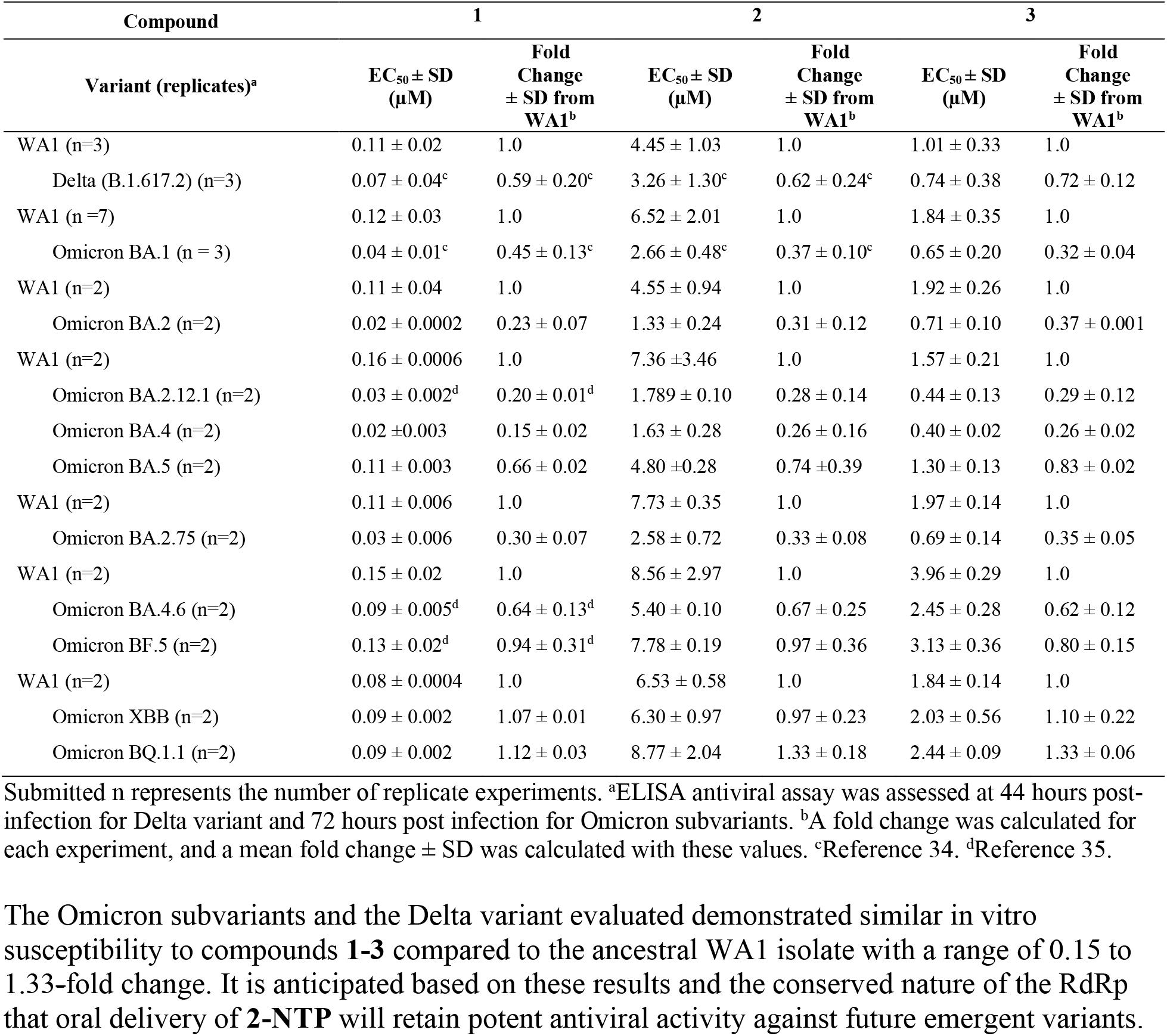
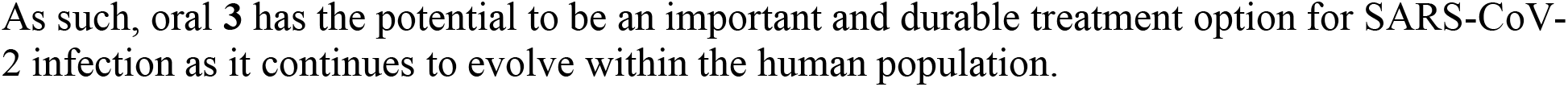
Antiviral activity of t**1**-**3** against SARS-CoV-2 variants.

Ester prodrugs are typically broken down rapidly by carboxyesterases that are expressed in the intestine and the liver. Prodrug **3** was evaluated as a substrate for three human carboxyesterases and found to be an excellent substrate for each with a short half-life of <4.3 min (Table 1). Consistent with the enzyme data, rapid hydrolysis was observed for **3** and the tri-ester prodrugs **5** and **6** in intestinal and hepatic S9 fractions. Mono-acetyl ester **5** was notably more stable in all species except rat and this is presumed to be due to its low log D that likely reduces its ability to be recognized by carboxyesterases. The plasma stability across the higher order species was generally greater than the intestinal and hepatic S9 stability for all the prodrugs. The combined in vitro stability results across these matrices suggest that **3** will be efficiently cleaved pre-systemically following absorption in vivo. A potential route of non-productive metabolism during absorption is oxidation of the adenosine-like *C*-nucleobase by the action of adenosine deaminase that is highly expressed in intestinal mucosa. Indeed, ester prodrugs of adenosine analogs have been previously designed to potentially alleviate adenosine deaminase metabolism during absorption.^36–37^ The metabolism of **2** and **3** by adenosine deaminase was therefore investigated in vitro and both compounds were found to be weak substrates or inhibitors of adenosine deaminase (see supporting information). The in-vitro data suggests that the oral bioavailability of **2** may not be significantly impacted by undesired intestinal metabolism.

To evaluate the impact of permeability on oral bioavailability, nucleoside **2** and prodrug **3** were initially dosed as solutions in rat, dog, cynomolgus and AGM (Table 2). The bioavailability of **2** in systemic circulation following oral administration of **3** was calculated based on the exposure of **2** determined from IV dosing normalized to the same mg-equivalent/kg of **2** dosed. Consistent with the in vitro stability data, only transient and low levels of intact prodrug **3,** when detectable, were observed at the early timepoints across all species. This was consistent with the in vitro stability observations in the intestinal and hepatic S9 fractions and established that the lung tissues in vivo are not exposed to any appreciable concentrations of the intact prodrug but only to the parent metabolite **2**. Given the relatively modest Caco-2 permeability and efflux improvements for **3**, we were surprised to observe a significant 2-, 3.6-and 7-fold improvement in the oral exposure of metabolite **2** in rat, AGM, and cynomolgus monkey, respectively. Indeed, the bioavailability of **2** that was achieved from oral dosing of **3** in cynomolgus monkey was comparable to the value reported for the highly lipophilic tri-isobutyryl ester prodrug **6** (F% = 28).^12^ Interestingly, similar ester-prodrug studies on the 4’-azido cytidine ribonucleoside analog, R1479, in cynomolgus monkeys concluded that a log P >2.0 was generally required for improved oral bioavailability in monkeys, and that less lipophilic 5’-monoesters with log P values below 2.0 were not very effective.^29^ The significantly improved oral properties of the adenosine analog **3** in cynomolgus monkey despite its log D < 2.0 may be the result of favorable interactions with intestinal nucleoside transporters which requires further investigation in the future. In dog, the bioavailability of **2** is very high and this was maintained by the prodrug **3**. Thus, the bioavailability of **2** following oral solution dosing of **3** was improved across multiple preclinical species tested and averaged 64% across rat, dog and cynomolgus monkeys.

For rapid clinical development, a stable crystalline form formulated into tablets was highly preferred. Therefore, the crystalline salt-free Form III of **3** that was identified from crystallization studies was dosed in dogs and cynomolgus monkeys in tablet form. A less than 2-fold drop to 67% oral bioavailability was observed in dogs, together with a 2-fold drop in oral bioavailability in cynomolgus monkey. In rats, suspension dosing was used and showed high oral bioavailability consistent with solution dosing. Despite the lower bioavailability of **2** in cynomolgus monkey, oral bioavailability was still 3.3-fold superior to the solution dosing of **2** leading to an average oral bioavailability from tablet dosing in dog and cynomolgus monkey of 42%. The favorable balance of permeability and solubility for **3** therefore resulted in improved oral delivery of **2** in both solution and solid forms. Taken together, these data supported the selection of **3** as the oral candidate.

The in vivo metabolism of **2** to **2-NTP** has been reported in lung tissue at 24 h following IV dosing in cyno, AGM, and ferrets.^11, 24^, ^27^ In these lung samples, **2-NTP** formation has been shown to be 10-to 15-fold less efficient from equimolar doses of **2** compared to the phosphoramidate **1,** consistent with the in vitro metabolism data described earlier. Therefore, both in vitro and in vivo data from IV administration supported the need for high systemic exposures of **2** to drive sufficient **2-NTP** formation in lung cells. Oral dosing results in a lower C_max_ compared to IV administration and it was not understood from the current data how the in vivo **2-NTP** concentrations in lung would be impacted by the different plasma-time exposure profile of **2** when delivered orally using the prodrug. Therefore, oral dosing of **3** was performed at 60 mg/kg in AGM to afford comparable AUC_0-24h_ exposures of **2** to those generated in the IV NHP experiments at 20 mg/kg reported in Table 2, and lung tissue samples were collected at 24 h for **2-NTP** analysis (Table 2). When normalized to the equivalent AUC_0-24h_ exposure of **2,** the data shows that **2-NTP** formed from oral administration is comparable to that of IV administration (Figure 2D) suggesting that lung **2-NTP** formation is more closely correlated with systemic AUC_0-24h_ exposures of **2,** rather than C_max_ concentrations. Given this observation, it is also reasonable to expect that antiviral efficacy is also correlated with the systemic exposures of **2** rather than C_max_ concentrations.

Recently, we have reported on the efficacy of **1** compared to IV administered **2** in a SARS-CoV-2 infection model in AGM and observed strong efficacy for the nucleoside **2** at high systemic exposures. ^24^ The same AGM model was then utilized to determine if efficacious concentrations of **2-NTP** in lung tissue could be achieved from systemic exposure to **2** through orally administered **3.** Oral treatment with **3** was initiated at 8 h post-infection with 60 or 120 mg/kg doses and then continued daily for 5 days. The doses were selected to provide a daily systemic exposure of **2** that bracketed the exposure from IV administered **2** at 20 mg/kg, which demonstrated strong efficacy relative to **1** in the earlier study.^24^ The viral loads (genomic RNA and infectious virus titer) were evaluated on bronchoalveolar lavage fluid (BALF), nasal, and throat swab samples, collected on 1, 2, 4, and 6 days post-infection (dpi) (Figure 3).

**Figure 3.**
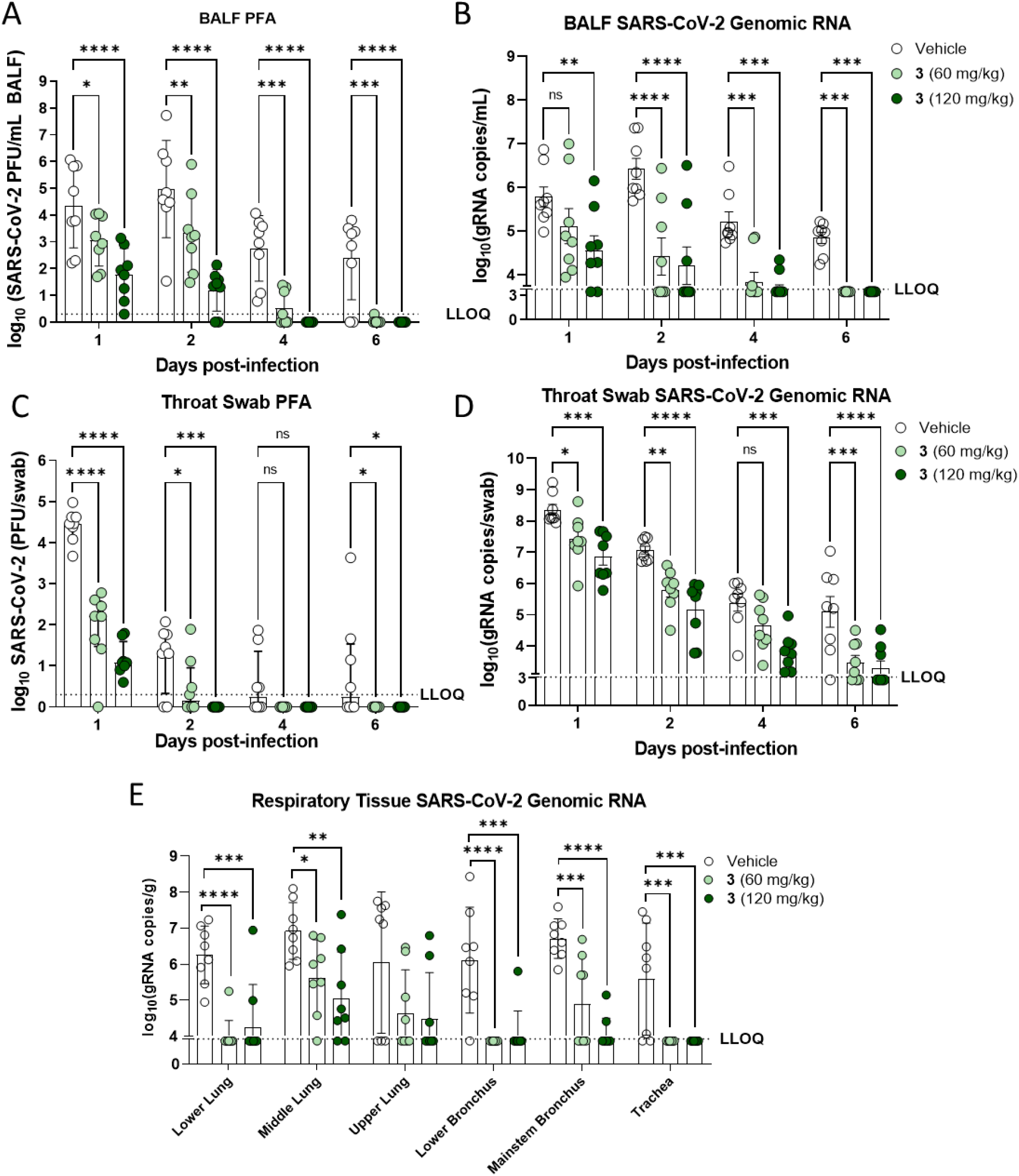
Antiviral effect of oral **3** on respiratory tract lavage, swab, and tissue samples in the African green monkey SARS-CoV-2 infection model. Panel **A**, Infectious virus in bronchoalveolar lavage fluid (BALF); Panel **B**, Genomic RNA in BALF; Panel **C**, Infectious viral titer in throat swab; Panel **D**, Genomic RNA in throat swab; Panel **E**, genomic RNA in respiratory tissue samples harvested day 6 across the respiratory tract. LLOQ, lower limit of quantification. *p<0.05; **p<0.01; ***P<0.001; ****p<0.0001.

At the end of the study, terminal respiratory tissues were harvested for evaluation of tissue viral RNA loads. Treatment of infected AGMs with either 60 or 120 mg/kg **3** resulted in a statistically significant decreases in infectious viral loads in the BALF throughout infection (Figure 3A). Treatment with **3** resulted in only 1 animal (in the 60 mg/kg group) with infectious SARS-CoV-2 in the BALF at or above the limit of quantification at day 6 post-infection, meanwhile 6 of the 8 vehicle animals had quantifiable infectious viral loads at this time-point (Figure 3A). The extent of infectious viral load reductions was dose-dependent, with larger reductions of infectious viral load observed for the group receiving **3** at 120 mg/kg over the sampling period. In line with reductions in infectious viral loads in the BALF, genomic RNA loads in BALF of **3** treated animals were also found to be significantly reduced at nearly all study timepoints assessed, with the one exception being the 60 mg/kg **3** treatment group at day 1 post-infection (Figure 3B). The results from RNA and infectious viral loads in the BALF highlights the significant effect of treatment with **3** in the lower airway.

In the upper airway, infectious viral titers and RNA were quantified from throat (Figure 3C-D) and nasal swabs (Supporting Information). In throat swabs, significant reductions in SARS-CoV-2 infectious (Figure 3C) and RNA loads (Figure 3D) were observed from the throat swabs at 1, 2, and 6 dpi from both dose groups. Furthermore, animals in the 120 mg/kg group showed additional reductions in viral RNA at day 4 post-infection. Notably, in both throat and nasal swabs, no detectable infectious virus was observed for any animal in either group receiving **3** after day 4 post-infection. There was a large observed difference in the SARS-CoV-2 genomic RNA levels and the infectious viral loads from the nasal swab analysis (Supporting Information). Vehicle control animals at day 1 post infection had a higher viral RNA load than that observed in BALF, but only half of the vehicle control animals had detectable levels of infectious virus. This difference highlights that viral RNA levels do not always correlate with infectious virus, and that viral RNA levels in the nasal cavity may, in part, be due to non-infectious viral RNA released from dead or dying cells. At the end of the study day 6, terminal respiratory tissues were also assessed for SARS-CoV-2 RNA loads, both doses resulted in significant reductions of viral RNA levels in 5 out of the 6 respiratory tissues evaluated (Figure 3E). Collectively, these data demonstrate that oral **3** at 60 mg/kg and 120 mg/kg are both highly efficacious at reducing infectious virus and viral RNA loads throughout the upper and lower respiratory tract of AGMs and similarly efficacious to **1**.^24^

Having established efficacy from oral dosing in AGMs we considered various methods to estimate the human efficacious dose of **3** for the clinic. One human dose estimation method employed C_max_/C_trough_ exposures of **2** relative to the in vitro SARS-CoV-2 antiviral EC_50_ potency of ∼ 1-2 µM in Vero E6 cells and primary lung cultures.^38^ However, a limitation of this method is that the relationship of plasma concentrations and in vitro antiviral EC_50_ has not been clearly established for SARS-CoV-2 efficacy in the clinic. Consideration was also given to a lung pharmacokinetic model that targeted doses of **3** that would generate equivalent lung **2-NTP** concentrations to that of IV administered **1** at its clinically efficacious dose. This resulted in an unfeasible multi-gram predicted dose of **3** and was inconsistent with the AGM efficacy study data. The 60 and 120 mg/kg doses in the AGM model were both highly efficacious despite the lung **2-NTP** concentrations at 24h being several-fold lower than the levels reported following IV administration of **1**.^12, 24^ A potential reason for this disconnect is the **2-NTP** concentrations from the gross tissue analysis at a single timepoint do not accurately reflect concentrations and distribution of **2-NTP** across infected cell-types in the lung. This is consistent with the different activation pathways of **3** and **1** to the common monophosphate metabolite which are likely cell-dependent and impacted by differences in the expression levels of the key enzymes.^39^ This is nicely illustrated by the fact that **1** is not effectively metabolized in Vero E6 cells, for example.^26^ Given the limited lung **2-NTP** data and the challenges in using this parameter to estimate efficacy we elected to use the plasma exposures of **2 (**AUC_0-24h_) and correlate this PK parameter to in vivo SARS-CoV-2 efficacy. In constructing an exposure-efficacy relationship model we combined all the efficacy data generated from the SARS-CoV-2 model in AGM for both oral **3** and IV **2**, in addition to the efficacy data reported in the ferret and mouse SARS-CoV-2 models using prodrug **6** (Table 5).^24, 27, 40^

**Table 5.** Plasma **2** Exposures and SARS-CoV-2 Antiviral Efficacy in Vivo.

| Cpd. | SARS-Cov-2 Model | Dose (mg/kg) | Route, regimen | **2**<br>C<sub>max</sub><br>(μM) | **2**<br>AUC<sub>0-24h</sub><br>(μM.h) | Log<sub>10</sub><br>gRNA<br>decrease<sup>a</sup> | Reference |
| --- | --- | --- | --- | --- | --- | --- | --- |
| **1** | AGM | 10 | IV, once daily | 12.6 | 7.6 | -1.38 | 24 |
| **2** | AGM | 7.5 | IV, once daily | ~33<sup>b</sup> | 55 | -1.02 | 24 |
| **2** | AGM | 20 | IV, once daily | 87 | 147 | -1.79 | 24 |
| **3** | AGM | 60 | Oral, once daily | 21 | 111<sup>c</sup> | -1.41 | This ref. |
| **6** | Ferret | 10 | Oral, twice daily | 5 | 54 | -1.98<sup>d</sup> | 27 |
| **6** | Mouse | 30 | Oral, twice daily | 13 | 95 | -3.29<sup>e</sup> | 40 |

<sup>a</sup>Decrease in genomicRNA (gRNA) relative to vehicle control in bronchioalveolar lavage sample on day 6 post-infection unless noted otherwise. <sup>b</sup>Estimated based on dose-proportionality to PK results obtained at 20 mg/kg.
<sup>c</sup>Calculated AUC<sub>0-24h</sub> at 60 mg/kg from the average of 14.4, 60 and 86.5 mg/kg AGM PO experiments in Table 2
<sup>d</sup>Nasal lavage sample on day 4 post-infection. <sup>e</sup>Lung tissue plaque forming unit (PFU) titer on final necropsy day 4 post-infection.

Including the data from the AGM IV study that evaluated both **1** and **2** in the same study was a critical aspect of the analysis because it is the only study, published up to now, that has directly compared the efficacy of **1** and **2** at clinically relevant exposures of **1**. Reported in Table 5 are the C_max_ and AUC_0-24h_ exposures of **2**, calculated using healthy animal PK, at the efficacious dose across each model. The efficacy in Table 5 from the ferret and mouse SARS-CoV-2 models is from the highest doses evaluated in those respective models following a treatment-based protocol.^27, 40^ Robust efficacy was achieved at plasma AUC_0-24h_ for **2** ranging from 54-111 µM.h. Thus, as anticipated, the systemic exposures of **2** that drive preclinical efficacy are >20-fold higher than the exposure of **1** at its 100 mg maintenance dose in human (AUC_0-24h_ = 2.6 µM.h).^24, 41^ Moreover, the efficacious exposure of **2** is also up to 17-fold higher than the exposure of **2** that is observed as a metabolite following the IV dosing of **1** (AUC_0-24h_ = 7.6 µM.h). It can be concluded that amidate prodrug **1** is very efficient at generating efficacious **2-NTP** concentrations in lungs from a relatively brief and low systemic exposure but has the disadvantage of IV administration. In contrast, **2** is much less efficient at metabolism to **2-NTP** so it requires a high systemic exposure and consequently higher doses for pre-clinical efficacy, but these exposures can be achieved orally using prodrug **3**.

Scaling of the preclinical plasma PK parameters to human using allometry and employing an average ∼42% oral bioavailability of **2** (average F% from the tablet dosing in dog and cynomolgus monkey), resulted in a projected twice-daily dose of **3** of 350-400 mg. This dose range is estimated to result in a human exposure of GS-441524 that matches the efficacious exposure range in the mouse and AGM models (AUC_0-24h_ = 95-111 µM.h). The dose is considerably higher on a per mole basis than the 100 mg maintenance dose of **1** which effectively reduces any benefit of the more facile synthesis of **3** due to its lower synthetic complexity. Nevertheless, the dose is attainable and would be considerably higher for oral **2** without the benefit of the prodrug improvements. The simulated human PK at this dose affords a C_max_ ∼ 7.5 µM and C_ave_ ∼4.6 µM, several-fold higher than the antiviral EC_50_ range of 2.5-3.3 µM reported for **2** in Table 3. We were delighted to find that the observed Phase 1 PK was in good agreement with the estimated human PK from the preclinical species.^42^ Based on the human phase 1 PK and safety data, a 350 mg twice-daily dose for 5 days is currently being assessed in two Phase 3 clinical trials (ClinicalTrials.gov Identifiers: NCT05603143 and NCT05715528) in people with or without risk factors for progression to severe disease.

## Conclusion

The development of convenient oral COVID-19 treatment options is critically important as SARS-CoV-2 infections persist and new variants can emerge. Consequently, a great deal of discussion has emerged in the public arena regarding the oral development of **2**, the parent nucleoside of **1**. However, our previous work on **2** had rigorously established significant hurdles for oral **2** existed, centered on its low bioavailability in non-human primates and the likely requirement for high systemic exposures to overcome its inefficient metabolism to **2-NTP**. A prodrug approach was therefore employed to lower the oral dose, reduce drug mass, and avoid loss of precious unabsorbed drug leading to the selection of a 5’-isobutyrl ester prodrug **3.** The ester prodrug **3** balances solubility, stability, and permeability and efficiently breaks down pre-systemically following oral delivery to provide high plasma concentrations of **2**. Despite only a modest improvement in permeability relative to **2,** improved oral bioavailability of **2** was found across multiple species following solution dosing and most importantly, solid tablet dosing. Oral administration of **3** was shown to generate the same common active **2-NTP** metabolite in the lungs of AGMs as **1** and drive a comparable antiviral effect to **1** in the AGM SARS-CoV-2 infection model. An efficacy-exposure relationship model was then constructed to estimate a human daily dose in the range of 350-400 mg twice daily. We also confirmed in vitro the potent activity of **1**, **2**, and **3**, against recent SARS-CoV-2 variants of concern. Oral administration of prodrug **3** effectively delivers the same active **2-NTP** metabolite as intravenous **1** and is currently being evaluated at a dose of 350 mg twice daily in two global Phase 3 COVID-19 trials. Further, the broad-activity established for **2-NTP** toward many RNA viruses also supports the potential to deploy **3** in a rapid-response scenario to combat future emerging RNA viruses with pandemic potential.

## EXPERIMENTAL SECTION

All organic compounds were synthesized at Gilead Sciences, Inc (Foster City, CA) unless otherwise noted. Commercially available solvents and reagents were used as received without further purification. Nuclear magnetic resonance (NMR) spectra were recorded on a Bruker 400 MHz at rt, with tetramethyl silane as an internal standard. Proton nuclear magnetic resonance spectra are reported in parts per million (ppm) on the *δ* scale and are referenced from the residual protium in the NMR solvent (CHCl_3_-*d_1_*: *δ* 7.26, MeOH-*d_4_*: *δ* 3.31, DMSO-*d_6_*: *δ* 2.50, CH_3_CN-*d_3_*: *δ* 1.93). Data is reported as follows: chemical shift [multiplicity (s = singlet, d = doublet, t = triplet), coupling constants (*J*) in Hertz. LCMS was conducted on an Agilent 1260 Infinity G6125B LCMS equipped with a Phenomenex Kinetex 2.6 µm C18 110 Å column (50 mm x 2.1 mm) eluting with 0.1% acetic acid in water (Buffer A) and 0.1% acetic acid in CH_3_CN (Buffer B); Gradient, 0-1.0 min 10-100% buffer B, 1.0-1.35 min 100% buffer B, then 1.35-1.36 min 100-10% CH_3_CN at 1.0 mL/min. Preparative normal phase silica gel chromatography was carried out using a Teledyne ISCO Combi Flash Companion instrument with silica gel cartridges. Purities of the final compounds were determined by high-performance liquid chromatography (HPLC) and were greater than 95%. HPLC purity was determined on an Agilent 1100 Series HPLC equipped with a Gemini column 2.6 µm 100Å column (100 mm × 4.6 mm) eluting with a 2-98% gradient of 0.1% trifluoroacetic acid in water and 0.1% trifluoroacetic acid in acetonitrile at a flow rate of 1.5 mL/min.

Compounds **1**, **2**, **6** and **7** have been reported previously.^12^

### Synthesis of ((2*R*,3*S*,4*R*,5*R*)-5-(4-aminopyrrolo[2,1-*f*][1,2,4]triazin-7-yl)-5-cyano-3,4-dihydroxytetrahydrofuran-2-yl)methyl isobutyrate (3)

To a solution of (3a*R*,4*R*,6*R*,6a*R*)-4-(4-aminopyrrolo[2,1-*f*][1,2,4]triazin-7-yl)-6-(hydroxymethyl)-2,2-dimethyltetrahydrofuro[3,4-*d*][1,3]dioxole-4-carbonitrile **4** (2000 mg, 6.0 mmol)^30^ and isobutyric acid (638 mg, 7.2 mmol) in DMF (5 mL), *N, N’*-diisopropylcarbodiimide (914 mg, 7.2 mmol) was slowly added followed by 4-dimethylaminopyridine (737 mg, 6.0 mmol). The reaction mixture was stirred for 4 h and then diluted with ethyl acetate, washed with water, brine, dried over sodium sulphate, and concentrated under reduced pressure. The residue was subjected to silica gel chromatography eluting with 20% MeOH in CH_2_Cl_2_ to provide the intermediate ((3a*R*,4*R*,6*R*,6a*R*)-6-(4-aminopyrrolo[2,1-*f*][1,2,4]triazin-7-yl)-6-cyano-2,2-dimethyltetrahydrofuro[3,4-*d*][1,3]dioxol-4-yl)methyl isobutyrate. LCMS *m/z* = 402.2 (M+1). To a solution of the intermediate (1500 mg) in THF (10 mL) conc. HCl (2 mL) was added, and the mixture stirred at r.t for 4 h. The reaction mixture was diluted with CH_2_Cl_2_, washed with water, saturated aqueous bicarbonate, and brine, dried over sodium sulphate, concentrated subjected to silica gel chromatography eluting with 30% MeOH in CH_2_Cl_2_ to afford the title compound (660 mg, 50%). ^1^H NMR (400 MHz, MeOH-*d*_4_) *δ* 7.88 (s, 1H), 6.96 – 6.85 (m, 2H), 4.50 – 4.27 (m, 4H), 4.16 (dd, *J* = 6.2, 5.3 Hz, 1H), 2.56 (p, *J* = 7.0 Hz, 1H), 1.14 (dd, *J* = 7.0, 3.8 Hz, 6H). LCMS m/z: 362.1 (M+1). ^13^C NMR (400 MHz, CHCl_3_-*d*_3_) *δ* 175.9, 155.6, 147.9, 123.5, 110.2, 100.8, 116.9, 116.6, 81.3, 79.0, 74.0, 70.2, 62.9, 33.2, 18.7, 18.6. HRMS m/z: 362.14615, C_16_H_20_N_5_O_5_ requires 362.14644.

### ((2*R*,3*S*,4*R*,5*R*)-5-(4-aminopyrrolo[2,1-*f*][1,2,4]triazin-7-yl)-5-cyano-3,4-dihydroxytetrahydrofuran-2-yl)methyl acetate (5)

The title compound was prepared from **4** (100 mg, 0.3 mmol) using the same method described above except acetic acid was used instead of isobutyric acid, and acetonitrile instead of THF in the acid deprotection step, to yield the title product (40 % overall yield). ^1^H NMR (400 MHz, DMSO-*d*_6_) δ 8.03 –7.96 (m, 3H), 6.92 (d, *J* = 4.5 Hz, 1H), 6.81 (d, *J* = 4.5 Hz, 1H), 6.31 (d, *J* = 6.0 Hz, 1H), 5.39 (d, *J* = 5.9 Hz, 1H), 4.70 (t, *J* = 5.5 Hz, 1H), 4.33 (dd, *J* = 11.9, 2.8 Hz, 1H), 4.23 (m, 1H), 4.14 (dd, *J* = 12.0, 5.9 Hz, 1H), 3.94 (q, *J* = 5.9 Hz, 1H), 2.02 (s, 3H). LCMS *m/z* = 334.1 (M+1).

### SARS-CoV-2-NLuc A549-hACE2 assay

Tested compounds are prepared in 100% DMSO in 384-well polypropylene plates (Greiner, Monroe, NC, Cat# 784201) as 4 replicates of 10 serially diluted concentrations (1:3). The serially diluted compounds were transferred to low dead volume Echo plates (Labcyte, Sunnyvale, CA, Cat# LP-0200). The test compounds were then spotted to 384-well assay plates (Greiner, Monroe, NC, Cat# 781091) at 200 nL per well using an Echo acoustic dispenser (Labcyte, Sunnyvale, CA). A549-hACE2 cells were harvested and suspended in DMEM (supplemented with 2% FBS and 1X Penicillin-Streptomycin-L-Glutamine) and seeded to the pre-spotted assay plates at 10,000 cells per well in 30 µL. SARS-CoV-2-NLuc virus^43^ was diluted in DMEM (supplemented with 2% FBS and 1X Penicillin-Streptomycin-L-Glutamine) at 350,000 plaque forming units (PFU) per mL and 10 µL per well was added to the assay plates containing cells and compounds (MOI 0.35). The assay plates were incubated for 2 days at 37°C and 5% CO2. At the end of incubation, Nano-Glo reagent (Promega, Madison, WI, Cat # N1150) was prepared. The assay plates and Nano-Glo reagent were equilibrated to room temperature for at least 30 min. 40 µL per well of Nano-Glo reagent was added and the plates were incubated at room temperature for 30 min before reading the luminescence signal on an EnVision multimode plate reader (PerkinElmer, Waltham, MA). Compound **1** was used as positive control and DMSO was used as negative control. Values were normalized to the positive and negative controls (as 0% and 100% replication, respectively) and data was fitted using non-linear regression analysis by Gilead’s dose response tool. The EC_50_ value for each compound was defined as the concentration reducing viral replication by 50%.

### SARS-CoV-2 NHBE cell assay

Compounds were evaluated for SARS-CoV-2 potency as described previously.^27^

### SARS-CoV-2 Nucleoprotein ELISA in A549-hACE2-TMPRSS2 cells

SARS-CoV-2 isolates for the ELISA antiviral assays were acquired through the World Reference Center for Emerging Viruses and Arboviruses at UTMB (Delta) and BEI Resources, National Institute of Allergy and Infectious Diseases (NIAID), National Institutes of Health (NIH). Isolates obtained from BEI Resources were deposited by the CDC (WA1 reference). Omicron variants obtained from BEI were deposited by Viviana Simon (BA.2.12.1), Andrew S. Pekosz (BA.4.6, BF.5, BQ.1.1) and Mehul Suthar (XBB). Omicron isolates BA.2, BA.4, BA.5, and BA.2.75 were obtained from the Gale laboratory, University of Washington. A total of 3×10^4^ A549-hACE2-TMPRSS2 cells in 100 μL DMEM (supplemented with 10% FBS and 1x penicillin-streptomycin) were seeded into each well of a 96-well plate and incubated overnight. The following day, medium was aspirated, and 100 μL of DMEM containing 2% FBS was added to each well. Three-fold serial dilutions of compound (in triplicate) were added to each well using an HP D300e digital dispenser with a final volume of 200 μL per well. Immediately after compound addition, cells were infected with 1.5 x 10^3^ PFU of the relevant SARS-CoV-2 variant (diluted in 100 μL of DMEM supplemented with 2% FBS), resulting in MOI 0.05. Plates were centrifuged for 2 min at 200 x g and then incubated at 37°C with 5% CO_2_ for 48 h (or 72 h for Omicron strains and the WA1 reference), after which medium was aspirated and cells were fixed with 100% MeOH for 10 min at rt. The MeOH was removed, and plates were air dried for 10 min at RT, followed by 1 h of incubation with 100 μL per well of blocking buffer (phosphate-buffered saline (PBS) with 10% FBS, 5% nonfat dry milk, and 0.1% Tween 20) for 1 h at 37°C. The blocking buffer was then aspirated, and 50 μL of a 1:4,000 dilution of rabbit anti-SARS-CoV-2 nucleocapsid antibody (Invitrogen, Waltham, MA, Cat# MA536086) in blocking buffer was added and incubated for 2 h at 37°C. Plates were washed 5 times with 100 μL per well of PBS containing 0.1% Tween 20 prior to addition of 50 μL per well of horseradish peroxidase-conjugated goat anti-rabbit IgG (ImmunoReagents, Raleigh, NC, Cat# GtxRb-003-FHRPX) diluted 1:4,000 in blocking buffer. Plates were again incubated for 1 h at 37°C and then washed 5 times with 100 μL PBS with 0.1% Tween 20. One hundred microliters 3,3′,5,5′-tetramethylbenzidene reagent (Thermo Scientific, Waltham, MA, Cat# ENN301) was added to each well and allowed to incubate at rt until visible staining of the positive-control wells, usually 5 to 10 min. The reaction was stopped with addition of 100 μL per well of 3,3′,5,5′-tetramethylbenzidene stop solution (SeraCare, Milford, MA, Cat# 5150-0021). The absorbance was then read at 450 nm using an EnVision plate reader. Fold change for variants was calculated for each experiment, with comparison to the relevant WA1 reference. Fold change across from each replicate for all experiments was then averaged to obtain the final reported values. EC_50_ is defined as the compound concentration at which there was a 50% nucleoprotein expression relative to infected cells with DMSO alone (0% inhibition) and uninfected control cells (100% inhibition). EC_50_ values were determined using GraphPad Prism 8.1.2 with nonlinear regression curve fits. Constraints were used when required to ensure the bottom or top of the fit curves were close to 0 and 100, respectively.

### Cytotoxicity Assay

Compounds (200 nL) were spotted onto 384-well black Greiner plates prior to seeding 5000 either A549-hACE2 or NHBE cells per well of in a volume of 40 µL culture medium. The plates were incubated at 37 °C for 48 h with 5% CO_2_. On day 2, 40 µL of CellTiter-Glo (Promega) was added and mixed 5 times. Plates were read for luminescence on an EnVision and the CC_50_ (compound concentration for reducing 50% of luminescence signal as a measure of cell viability) were calculated using a nonlinear regression model (four parameters).

The 5-day CC_50_ assays for MRC5 and NHBE are described in the Supporting Information. The 5-day CC_50_ assays for PC-3, MT-4, quiescent PBMC, stimulated PBMC, and freshly isolated PHH are conducted as reported.^32^

### In vitro metabolism

A549-hACE2 or NHBE cells were seeded in a 12-well plate at 2.5×10^5^ cells per well. 24 h later, cell-culture medium was replaced with medium containing compound, as indicated, and incubated at 37 °C with 5% CO_2_. At the indicated time post compound addition, cells were washed 3 times with ice-cold tris-buffered saline, scraped into 0.5 mL ice-cold 70% MeOH and stored at –80 °C. Extracts were centrifuged at 15,000 g for 15 min and supernatants were transferred to clean tubes for evaporation in a miVac Duo concentrator (Genevac, Painter, NY). Dried samples were reconstituted in mobile phase A containing 3 mM ammonium formate (pH 5) with 10 mM dimethylhexylamine in water for analysis by LC-MS/MS, using a multi-stage linear gradient from 10% to 50% acetonitrile in mobile phase A at a flow rate of 360 μL/min. Analytes were separated using a 50 x 2 mm, 2.5 μm Luna C18(2) HST column (Phenomenex, Torrance, CA) connected to an LC-20ADXR (Shimadzu, Kyoto, Japan) ternary pump system and an HTS PAL autosampler (LEAP Technologies, Carrboro, NC). Detection was performed on a Qtrap 6500+ (AB Sciex, Redwood City, CA) mass spectrometer operating in positive ion and multiple reaction monitoring modes. Analytes were quantified using a 7-point standard curve ranging from 0.624 to 160 pmol per million cells prepared in extracts from untreated culture wells that were counted for each timepoint.

### Stability in pH2 and pH7 buffered solutions

The aqueous stability of compounds was assessed over a time of not less than 24 h at 40 °C. Stability was determined in 50 mM phosphate buffered solutions at both pH2 and pH7, with 150 mM NaCl. For each compound and pH condition, 50 µg/mL samples were prepared with not more than 50% acetonitrile as a cosolvent. Samples were analyzed by UPLC using a Waters Acquity UPLC with a PDA UV detector.

### Thermodynamic Solubility in pH 2 water

The aqueous solubility of compounds was assessed over a time of not less than 72 h. Solubility was determined at ambient temperature in water adjusted to pH 2 with 1N HCl. Solids were added to 0.5 mL of pH 2 water in 1.5-mL Eppendorf tubes, vortexed, then agitated for 24 h in an Eppendorf ThermoMixer C at 1400 rpm. To determine concentration in solution, the suspensions were centrifuged for 3 min at 15,000 rpm. Supernatants were diluted by a factor of 100 with 50:50 v/v acetonitrile:water. All diluted supernatants were analyzed by UPLC using a Waters Acquity UPLC with a PDA UV detector.

### Thermodynamic Solubility in pH7 buffered solution

The aqueous solubility of compounds was assessed over a time of 24 h. Solubility was determined at ambient temperature in a 50 mM phosphate buffered pH7 solution with 150 mM NaCl. Solids were added to the buffered solution in 1.5-mL Eppendorf tubes, vortexed for 1 min, then agitated for 24 h in an Eppendorf ThermoMixer C. To determine concentration in solution, the suspensions were centrifuged for 15 min at 15,000 rpm. Supernatants were diluted to a volume of 1 mL with 30:70 v/v acetonitrile:water. All diluted supernatants were analyzed by UPLC using a Waters Acquity UPLC with a PDA UV detector.

### Carboxyesterase (CES) stability

Test compounds or positive control substrates (oseltamivir for CES1 enzymes or procaine for CES2) were incubated with individual Supersome preparations (Corning Life Sciences, Corning, NY; final CES concentration 1.5 mg/mL) in 0.1 M potassium phosphate buffer (pH 7.4) at 37 °C. Substrates were added to a final concentration of 2 µM to initiate the reaction. The final incubation volume was 250 mL. Aliquots were removed after incubation for 0, 10, 30, 60 and 120 min. The reactions were stopped by the addition of quench solution (90/10 (v/v) CH_3_CN/MeOH with 0.1% formic acid) containing internal standard. Following protein precipitation and centrifugation, supernatant was diluted with an equal volume of water prior to analysis. For procaine, supernatant was dried down and reconstituted with water. All samples were analyzed by LCMS/MS and peak-area ratios were used for quantification. Analysis by LC-MS/MS was performed on a Thermo Q-Exactive mass spectrometer coupled to a Dionex UltiMate 3000 HPLC with a Leap Technologies HTC PAL autosampler. Separation of test compounds was accomplished using a Waters Acquity BEH C18, 50 x 2.1 mm, 1.7 µm column using a multi-stage linear gradient with mobile phases containing 0.1% formic acid in either water or acetonitrile, at a flow rate of 0.2 mL/min.

### Intestinal and Hepatic S9 Stability

Stability of the compounds was assessed in both intestinal and hepatic S9 fractions (BioIVT, Baltimore, MD) from select species. For S9 stability, duplicate aliquots of test compound or positive control substrate (GS-7340) were added to either PMSF-free intestinal or hepatic S9 stock diluted with 100 mM phosphate buffered saline, pH 7.4, to obtain a protein concentration of either 1.0 or 2.4 mg/mL, respectively. The S9 metabolic reactions were initiated by the addition of the substrates to the S9 reaction mixture to a final concentration of 2 µM. At 0, 10, 20, 30, 60 and 120 min (intestinal S9) or at 2, 12, 25, 45, 65 and 90 min (hepatic S9), 25 µL aliquots of the reaction mixture were transferred to plates containing 225 µL of internal standard in quenching solution (acetonitrile). After quenching, the plates were centrifuged at 3000 x g for 30 min, and 150 µL aliquots of each supernatant were diluted with 150 µL water. Aliquots (10 µL) of the diluted supernatant were analyzed by LC-MS/MS as described for CES stability assessment.

### Plasma Stability

Stability of the compounds was assessed in plasma from select species. Duplicate aliquots of plasma were warmed to 37 °C and the metabolic reactions initiated by the addition of test compound (6 µL of 0.1 mM DMSO stock) or plasma stability standard (GS-7340) to obtain a final substrate concentration of 2 µM. At 0.05, 0.5, 1, 2, 3 and 4 h plasma, 25 µL aliquots of the reaction mixture were treated and analyzed following the S9 stability method described above.

### Caco-2 Permeability

The bidirectional permeability of tested compounds was assessed in pre-plated Caco-2 cells (clone C2BBe1), obtained from Sigma-Aldrich, Inc. (Atlanta, GA). Cell monolayers were grown to confluence on collagen-coated, microporous, polycarbonate membranes in 24-well transwell plates for 21 days. The permeability assay buffer in donor wells was Hanks’ balanced salt solution (HBSS) containing 10 mM HEPES and 15 mM glucose at a pH of 6.5 containing 200 µM BNPP. The receiver wells used HBSS buffer containing 10 mM HEPES and 15 mM glucose at a pH of 7.4 and supplemented with 1% BSA. After an initial equilibration with transport buffer, TEER values were read to test membrane integrity. The experiment was started by the addition of buffers containing test compounds, 200 µL and 1000 µL in the apical and basolateral chamber, respectively, to determine forward (A to B) and reverse (B to A) permeability. At 0 and 2 h post dose, 10 µL was sampled from donor compartment and was diluted in 190 µL of 20% MeOH. At 1 and 2 h post dose, 100 µL of solution was taken from the receiver compartments and was diluted in 100 µL of 20% MeOH. Removed buffer was replaced with fresh buffer and a correction was applied to all calculations for the removed material. Each compound was tested in 2 separate, replicate wells for each condition. All samples were then extracted with 400 µL of 100% acetonitrile containing internal standard, to precipitate protein. To test for non-specific binding and compound instability, the total amount of drug was quantitated at the end of the experiment and compared to the material present in the original dosing solution as a percent recovery. Samples were analyzed by LC-MS/MS for quantitation of both prodrug and parent in each chamber on an API 6500+ MS system (Sciex, Framingham, MA) with Shimadzu LC-20ADXR ternary pump system (Shimadzu, Columbia, MD) and an HTC PAL autosampler from LEAP Technologies (LEAP Technologies, Carrboro, NC). Compounds were separated and eluted on a Synergi 4 µm Polar-RP 2.0 × 150 mm column (Phenomenex, Torrance, CA) at a flow rate of 0.6 mL/min and using a multi-stage gradient with mobile phases containing either 1 or 99% CH_3_CN in 0.2% formic acid.

### In vivo pharmacokinetics

In vivo pharmacokinetic (PK) studies were performed at either Covance (rat, dog and cynomolgus monkey; Madison, Wisconsin) or Lovelace Biomedical Research Institute (AGM; LBRI, Albuquerque, NM) in accordance with local IACUC guidelines. Rats, dogs, cynomolgus or AGM received a single intravenous (**2** as a 30-min infusion) or oral (**2** or **3**) administration at doses indicated and in vehicles noted below. Formulation of **2** for IV administration in rat, cyno, and AGM was 5% ethanol (EtOH), 30% propylene glycol (PG), 45% polyethylene glycol-300 (PEG300) and 20% water, pH 2-3 or in dog was 5% EtOH, 30% PG, 45% polyethylene glycol-400 (PEG400) and 20% water, with 1 equivalent of hydrochloric acid (HCl). Formulation of **2** for oral dosing in rat was 5% EtOH, 30% PG, 45% PEG400, and 20% Water, plus 1 equiv. of HCl or 2.5% dimethyl sulfoxide (DMSO), 10% Kolliphor HS-15, 10% Labrasol, 2.5% PG and 75% water, pH 2. Formulation of **2** for oral dosing in dog and cynomolgus monkey was 5% EtOH, 30% PG, 45% PEG400 and 20% water, with 1 equivalent of HCl. Formulation of **2** for oral dosing in AGM was 5% EtOH, 45% PEG300, 30% PG and 20% water, pH 2-hand-compressed **3**. Formulation of **3** for oral dosing in rat, cynomolgus monkey and AGM was 2.5% DMSO, 10% Kolliphor HS-15, 10% Labrasol, 2.5% PG and 75% water, pH 2-3; Formulation of **3** for oral dosing in dog was 0.5% DMSO, 2% Kolliphor HS-15, 2% Labrasol, 0.5% PG and 95% water. Formulation of **3** for solid oral dosing in rat was 0.5% methyl cellulose, 99.5 % water, pH 6. Formulation of **3** for solid oral dosing in pentagastrin pre-treated dogs (at a 175 mg fixed dose) was a hand compressed tablet with a composition of 50% GS-5245 freebase Form III, 44.5% microcrystalline cellulose, 4% crospovidone, and 1.5% magnesium stearate. Formulation of **3** for solid oral dosing in cynomolgus monkey (at a 124 mg fixed dose) was a hand compressed tablet with a composition of 50.8% GS-5245 freebase Form III, 43.7% microcrystalline cellulose, 4% crospovidone, and 1.5% magnesium stearate.

Serial blood collection was performed from 3 animals and processed to plasma at IV predose, 0.25, 0.48, 0.58, 0.75, 1.5, 3, 6, 8, 12, 24 h or oral predose, 0.25, 0.5, 1, 2, 4, 6, 8, 12 and 24 h post-dose. 20-50 µL aliquots of plasma were added to a mixture containing 250 μL of MeOH and 25 μL of internal standard solution and centrifuged, and 250 μL of resulting supernatant was then transferred and dried under a stream of nitrogen at 40°C and reconstituted in a mixture of 5% acetonitrile and 95% water. Plasma concentrations of **3** and/or **2** were determined using 8-point calibration curves spanning at least 3 orders of magnitude with quality control samples to ensure accuracy and precision, all prepared in naïve plasma. Analytes were separated on a 50 × 3.0 mm, 2.5 μm Synergi Polar-RP column (Phenomenex, Torrance, CA) using a multi-stage linear gradient with mobile phases containing 0.1% formic acid in either water or acetonitrile, at a flow rate of 0.7 mL/min. Pharmacokinetic parameters were calculated using Phoenix (v8, Certara, Princeton, NJ).

AGM lung tissue concentrations were determined as previously described.^12^ Briefly, lung tissues were collected as a non-survival surgical procedure at 24 h post-dose. Animals were sedated with ketamine administered via an intramuscular injection prior to surgery. A section of the lower lung was dissected and immediately placed into liquid nitrogen <5 min from the start of surgery. Frozen tissues were pulverized using a cell crusher and transferred into pre-weighed conical tubes, all performed and maintained on dry ice. Tissues were weighed and a volume of dry ice-cold extraction buffer was added. Resulting mixtures were then promptly homogenized. Un-dosed control tissues were used for quantification of the metabolites in lung tissue, generated by spiking an appropriate amount of metabolite standard solution into control tissues. An aliquot of the homogenate was filtered, evaporated to dryness and reconstituted with 1 mM ammonium phosphate buffer for analysis by LCMS/MS. Analysis was performed using similar methods as previously described.^15^

### AGM SARS-CoV-2 antiviral efficacy study

The in vivo efficacy study was conducted at LBRI. All studies were conducted under an IACUC approved protocol in compliance with the Animal Welfare Act, PHS policy, and other federal statutes and regulations relating to animals and experiments involving research animals. The efficacy study, which involved animals experimentally infected with SARS-CoV-2 was conducted in an animal biosafety level 3 (ABSL-3) laboratory. Wild-caught AGM (St. Kitts origin) were sourced through Worldwide Primates Inc. (Florida, USA) for all studies. AGMs were housed in adjacent individual cages, within a climate-controlled room with a fixed light/dark cycle (12-h light/12-h dark). AGM were monitored at least twice daily for the duration of the study by trained personnel. Commercial monkey chow, treats and fruit were provided twice daily. Water was available to the AGM *ad libitum*. Antiviral efficacy of **3** was evaluated in 24 animals (12 males and 12 females) in 3 cohorts of 8 animals each staggered by 1 day. Animals were infected under anesthesia with 3×10^6^ TCID_50_ of SARS-CoV-2 virus (WA1 isolate) via intranasal (0.5 mL each nostril = 1 mL total) and intratracheal (2 mL) instillation. Animals were randomly placed into one of 3 groups (vehicle control, 60 mg/kg **3**, or 120 mg/kg **3**) each with an N=8. Per os (PO) dosing for all groups was initiated via oral gavage at 8 h post-infection and then once daily thereafter through day 5 post-infection. Compound **3** was formulated at either 30 mg/mL or 60 mg/mL solutions in 2.5% dimethyl sulfoxide (DMSO), 2.5% propylene glycol, 10% Labrasol, 10% Kolliphor HS-15, and 75% water at pH 2-3. Blood samples were collected from a femoral, saphenous, or cephalic vein daily into K_2_EDTA tubes. A daily blood samples was collected from each animal immediately prior to gavage to monitor dosing trough plasma PK levels. Samples were centrifuged at 1700-1800 ×*g* at 4 °C for 10 min, and plasma isolated, aliquoted, and frozen immediately on dry ice and stored at −70 °C until inactivation by organic solvent sterilization and subsequent bioanalysis. Nasal and throat swabs were collected at 1, 2, 4, and 6 days post-infection (dpi) using a cotton-tipped applicator presoaked in sterile saline. Swabs were placed in a tube containing 0.5 mL sterile saline, and frozen immediately on dry ice, and stored at −70 °C until further processing. Bronchioalveolar lavage fluid (BALF) was collected from left and right caudal lung lobes at 1, 2, 4, and 6 DPI. For BALF collection a pediatric bronchoscope (Olympus XP-40) was advanced into the caudal lung lobe, 10 mL of sterile saline was infused, and the maximum volume was aspirated. BALF was centrifuged at 1000 ×*g* for 10 min at 4 °C. The resulting cell pellet and 1 mL aliquots of supernatant were flash frozen and stored at −70 °C until further processing for viral titer analysis or inactivation by organic solvent sterilization and subsequent bioanalysis. Animals were monitored daily for any clinical evidence of disease and body weights and temperature measurements were recorded on day 1, 2, 4 and 6 post infection. Animals were euthanized for tissue collection and necropsy at day 6 post-infection. Sections of the right lung were flash frozen in liquid nitrogen and stored at −70 °C for analysis of viral RNA loads. Reverse transcription quantitative PCR (RT-qPCR) and plaque forming assay (PFA) methods are described in supporting information.

## ASSOCIATED CONTENT

### Supporting Information

The supporting information is available free of charge. Supporting information includes additional experimental details, supporting data tables and figures, NMR and HPLC spectral data (PDF) and molecular formula strings (CSV).

### Corresponding Author

. Richard L. Mackman, Gilead Sciences, Inc., Foster City, California 94404, United States.

### Current Addresses

^‡^Andrea Ambrosi: 1 DNA Way, South San Francisco, CA 94080, USA

### Author contributions

These authors contributed equally to the data generation and manuscript preparation; Darius Babusis and Jared Pitts.

### Notes

The authors declare the following competing interests. Some authors are current or former employees of Gilead Sciences and may own company stock.

## ACKNOWLEGDEMENTS

We would like to extend the following acknowledgments: Dr. Meghan Vermillion and the Lovelace Biomedical support team for execution of the SARS-CoV-1 AGM efficacy model. The following reagents were deposited by the Centers for Disease Control and Prevention and obtained through BEI Resources, NIAID, NIH: SARS-Related Coronavirus 2, Isolate hCoV-19/USA/NY-MSHSPSP-PV56475/2022 (Lineage BA.2.12.1; Omicron Variant), NR-56781, deposited by Dr. Viviana Simon; SARS-Related Coronavirus 2, Isolate hCoV-19/USA/MD-HP35538/2022 (Lineage BA.4.6; Omicron Variant), NR-58715 and SARS-Related Coronavirus 2, Isolate hCoV-19/USA/MD-HP34985/2022 (Lineage BF.5; Omicron Variant), NR-58716, contributed by Dr. Andrew S. Pekosz. We thank Kenneth S. Plante, Jessica A. Plante, and David S. Blakeman for coordination of virus stocks from the World Reference Center for Emerging Viruses and Arboviruses at the University of Texas Medical Branch.

## ABBREVIATIONS

AB: apical to basolateral
AGM: African green monkey
ATP: adenosine triphosphate
BALF: bronchioalveolar lavage fluid
BEGM: bronchial epithelial growth medium
CES: carboxyesterase
Cyno: cynomolgus
DIC: *N*,*N*′-Diisopropylcarbodiimide
DMAP: 4-dimethylaminopyridine
DMEM: Dulbecco’s modified eagle medium dpi, day post-infection
FBS: fetal bovine serum
GI: gasterointestinal
gRNA: genomic RNA
Hep: Hepatocytes hr, human recombinant
IACUC: International animal care and use committee
IV: intravenous
LLOQ: lower limit of quantification
MERS: Middle East Respiratory Syndrome
MOI: multiplicity of infection
NHBE: normal human bronchial epithelial
NHP: non-human primate
NTP: nucleotide triphosphate
PBMC: peripheral blood mononuclear cells
PBS: phosphate-buffered saline
PFA: Plaque forming assay
PFU: plaque forming unit
PHH: primary human hepatocyte
PMSF: phenylmethylsulfonyl fluoride
RdRp: RNA-dependent RNA polymerase
RSV: respiratory syncytial virus
SARS: severe acute respiratory syndrome
VOC: variant of concern
WHO: World Health Organization

## Table of Content Graphic

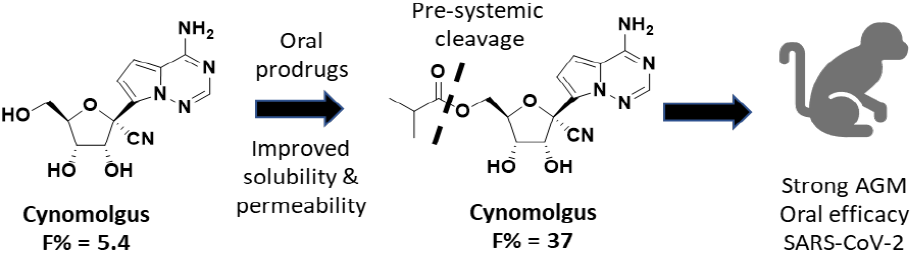

