## Supporting Information for "Discovery of GS-5245 (Obeldesivir), an Oral Prodrug of Nucleoside GS-441524 that Exhibits Antiviral Efficacy in SARS-CoV-2 Infected African Green Monkeys"

### Contents


| | |
|---|---|
| Cell Lines | S2 |
| Cytotoxicity Profiling Methods | S3 |
| SARS-CoV-2 African Green Monkey Study Assays | S4 |
| Adenosine Deaminase Profiling Methods | S6 |
| Crystalline Form Isolation and Characterization | S7 |
| Supporting Information (SI) Figures and Tables | S8 |
| References | S20 |
| Compound **3** and **5** Spectra | S21 |

### Cell Lines

The A549-hACE2 cell line was established and provided by the University of Texas Medical Branch (Mossel, **2005**) and were maintained in Dulbecco's Minimum Essential Medium (DMEM) (Corning, NY, Cat. #15-018CM) supplemented with 10% fetal bovine serum (FBS) (Hyclone, Logan, UT, Cat. #*SH30071-03*), 1X Penicillin-Streptomycin-L-Glutamine (Corning, Cat. #*30-009-CI*) and 10 µg/mL blasticidin (Life Technologies Corporation, Carlsbad, CA, Cat. #A11139-03). Cells were passaged 2 times per week to maintain sub-confluent densities. A549-hACE2-TMPRSS2 cells (Cat. #a549-hace2tpsa) were purchased from InvivoGen (San Diego, CA). Normal human bronchial epithelial (NHBE) cells were purchased from Lonza (Walkersville, MD) and maintained in bronchial epithelial cell growth medium (BEGM) (Lonza, Walkersville, MD) with all provided supplements in the BulletKit. Cells were passaged 1-2 times per week to maintain sub-confluent densities and were used for experiments at passages 2-4.

**Cytotoxicity Profiling Methods**

Sample compounds are serially diluted in quadruplicate 1:3 in ten points and are prespotted by acoustic transfer (ECHO) into replicate black polystyrene tissue culture-treated 384-well plates (Greiner 781946) for each adherent cell line in quadruplicate at 400 nL/well, and 310 nL/well for PBMCs. 1 mM puromycin is spotted at the edge as a positive control. Transfers are tracked with pipetting logs.

Cell lines are batch prepared and diluted to 16,600/mL to achieve a density of 1,500/well for galactose dependent-HepG2 and galactose dependent-PC-3, MRC-5, and 5600/mL for Huh7 to achieve 500/well in 90 µL of appropriate culture media. PBMCs are prepared at 75,000 cells/mL to obtain 5000 cells/well in 70 µL.

Each cell line is dispensed to their prespotted assay plates via µFlo to their designated volumes. The DMSO concentration in the final assay plates was 0.44% (v/v). Cells were incubated with compound for 5 days at 37°C in a $CO_2$ incubator. Puromycin (44 µM final concentration) and DMSO (0.44% v/v) were used on each assay plate as controls for 100% and 0% cytotoxicity, respectively. In addition, a $CC_{50}$ value for puromycin was determined for each cell line tested as a positive internal control and to assess assay sensitivity in each cell type.
At the end of the incubation period, processing was performed using a preprogrammed µFlo dispenser connected to an EL405 plate washer (Biotek). For adherent cells, in the first step, media from the 384-well assay plates was aspirated and cells were washed once with 80 µL Dulbecco’s phosphate buffered saline (PBS). In the next step, twenty microliters of CellTiter-Glo (Promega, Madison, WI) was added to each well of the plates using the µFlo. For PBMCs, media was aspirated down to 30 µL and an equivalent volume of Cell titer glo was added. Plates were a placed on a rotator and gently swirled at 300 rpm 20-30 minutes to mix.

Luminescence was measured with an EnVision plate reader (Perkin Elmer, Waltham, MA) and $CC_{50}$ values calculated from the data using standardized curve fitting programs.

### SARS-CoV-2 African Green Monkey Study Assays

**Reverse transcription quantitative PCR (RT-qPCR).** Tissue samples of ~100 mg each were homogenized in lysis buffer using a TissueLyser (Qiagen) and then centrifuged at 10,000 x g for 1 minute. The clarified supernatant was aliquoted into samples for RNA extraction. RNA was isolated from homogenized tissue supernatants, BALF and swab samples using the Direct-zol RNA purification kit (Zymo Research) following manufacturer's protocols. RT-qPCR reactions were set-up in triplicate using TaqMan Fast Virus 1-step Master Mix (ThermoFisher) with the following cycling conditions: 50°C – 5 minutes, 95°C – 20 seconds, and 40 cycles of 95°C for 3 seconds and 60°C for 30 seconds. The primer-probe set used for genomic RNA detection targeted the nucleocapsid gene (N gene) was as follows: N forward primer 5′ TTACAAACATTGGCCGCAAA 3′; N reverse primer 5 ′ GCGCGACATTCCGAAGAA 3′; N probe: 5' 6FAM-ACAATTTGCCCCCAGCGCTTCAG-BHQ-1 3'. The primer-probe set used for the sub-genomic RNA used a forward primer targeting the leader sequence in the 5'UTR and a reverse primer and probe targeting the envelope gene (E gene): sgLead SARS-CoV-2 Forward: 5′ CGATCTCTTGTAGATCTGTTCTC 3′; E Sarbeco Reverse: 5′ ATATTGCAGCAGTACGCACACA 3′; E Sarbeco Probe: 5' 6FAM-ACACTAGCCATCCTTACTGCGCTTCG-BHQ-1 3'. Genomic or sub-genomic copies were calculated using appropriate standard curves for each primer-probe set.

**Plaque forming assay (PFA).** Vero-TMPRSS2 cells expressing human transmembrane serine protease 2 (hTMPRSS2) were purchased from JCRB cell bank (Cat. #JCRB 1818), National Institutes of Biomedical Innovation, Health and Nutrition. Cells were maintained at 37°C and 5% $CO_2$ in Dulbecco's Minimum Essential Medium (DMEM) with GlutaMAX (Gibco Cat. #10569-010) supplemented with 10% heat-inactivated fetal bovine serum (FBS) (Hyclone Cat. #SH30396.03), 100 units/mL penicillin, 100 µg/mL streptomycin (Gibco Cat. #15140-122), and 1 mg/mL Geneticin. Cells were passaged 2-3 times per week with 0.25% Trypsin/0.02% EDTA (Gibco Cat. #25200056) and seeded for experimental set-ups between passage 10 and 30. $1\times10^6$ Vero-TMPRSS2 cells/well were seeded into 6-well plates in 2 mL of maintenance media and incubated overnight at 37°C and 5% $CO_2$. The following day, cells were visualized under a light microscope to confirm confluency of >95%. Samples were serially diluted 10-fold in infection medium (DMEM + 2% FBS) to a final dilution of $10^{-3}$ or $10^{-4}$. Spent supernatant from cultures was aspirated and replaced with 250 µL of serially diluted inoculum/well in duplicate, and culture plates were returned to the incubator for 1 h with gentle rocking every 15 min. Following incubation, 5 mL of pre-warmed overlay medium (DMEM with 2% FBS, 1X penicillin/streptomycin, and 1.5% carboxymethylcellulose) was added to each well. Cells were then incubated at 37ºC and 5% $CO_2$ without agitation for 3 days, at which point 5 mL of crystal violet fix/stain solution was added to each well. Cells were incubated at room temperature overnight. Supernatants containing the crystal violet solution were discarded, and wells were washed with water 2 to 4 times each until plaques were visible and washes were clear of crystal violet residue. Plaques were counted manually from the most dilute wells consistently containing >5 plaque forming units (PFU).

**Statistical analyses**. Statistics were performed using GraphPad Prism 8.0. In all analyses, compound **3** treatment groups were compared independently against the vehicle control group. For longitudinal analyses (BALF and swabs samples), results were analyzed by two-way ANOVA with Bonferroni post-hoc correction for multiple comparisons. For terminal tissues, the compound **3** treatment groups were individually compared to the vehicle control using one-way

ANOVA with Bonferroni post-hoc correction. Samples analyzed by RT-qPCR or plaque assay which were below the lower limit of quantification for the assay were assigned a value of 1/2 of the lower limit of quantification then log transformed. Corrected $p$ values of <0.05 were considered statistically significant.

**Adenosine Deaminase Profiling Methods**

**Effect of test articles on catalytic activity of recombinant human adenosine deaminase (ADA1).** Effect of test articles on catalytic activity of recombinant human ADA1 was evaluated by measuring conversion of adenosine to inosine by ADA1 in the presence and absence of test articles. 100 μM of each test article was mixed with 100 μM adenosine and reaction was initiated by addition of 3 nM recombinant human ADA1. Reaction was carried out at room temperature in 100 μL of buffer containing 20 mM HEPES, pH 8.0. All concentrations are final after mixing. Conversion of adenosine to inosine catalyzed by ADA1 in the absence (1% DMSO) and the presence of test article was monitored by measuring absorbance at 265 nm as function of time (SI Figure 4, SI Table 2a). Half-life ($t_{1/2}$) of the conversion was calculated by non-linear least squares data fit to a single exponential decay equation with GraphPad Prism software. Control reactions were also performed in the presence of 10 μM ADA1 inhibitor EHNA ($K_i$ = 4 nM).

**Evaluation of potential for deamination of test articles by recombinant human adenosine deaminase (ADA1).** Ability of a recombinant human ADA1 to catalyze deamination of a test article was evaluated by measuring absorbance of a test article during incubation with ADA1 for up to 90 min at room temperature. 100 μM test article was mixed with 3 nM recombinant human ADA1 in 100 μL total volume in a buffer containing 20 mM HEPES, pH 8.0 to initiate reaction. All concentrations are final after mixing. Absorbance spectra were collected from 230-300 nm. Adenosine and inosine at 100 μM were used as a positive and a negative control, respectively. Absorbance and a percent of absorbance change of adenosine (positive control) and inosine (negative control) after 90 min incubation with ADA1 were determined at 265 nm. Percent of absorbance change was calculated as 100%*(Absorbance at 0 min - Absorbance at 90 min)/(Absorbance at 0 min). Absorbance and absorbance change of test articles after 90 min incubation with ADA1 were determined at 245 nm and 285 nm and compared to absorbance of inosine at respective wavelengths (SI Figure 4, SI Table 2b). Wavelength at which absorbance for each molecule was measured was selected to observe maximal difference between absorbance of test article and inosine. Control reactions were also performed in the presence of 10 μM ADA1 inhibitor EHNA ($K_i$ = 4 nM).

**Crystalline Form Isolation and Characterization**

A representative procedure for the isolation of **3** crystalline freebase Form III is as follows:

Compound **3** (18.7 g) was dissolved in MeCN (140 mL) and the internal temperature was adjusted to 20°C. Concentrated aqueous HCl (3 mol equivalents) was added, and the reaction mixture was agitated until the reaction was deemed complete. The resulting solids were isolated by vacuum filtration and rinsed with MeCN. The isolated solids were charged back to the reactor and suspended in MeTHF (140 mL). The internal temperature was adjusted to 20°C and the slurry was washed with 15% aq. $KHCO_3$, and water. The solvent was exchanged to a mixture of MeCN (160 mL) and DCM (80 mL), and the resulting slurry was seeded with compound **3** Form III (0.5 wt%). The internal temperature of the slurry was adjusted to -10°C and aged. The solids were isolated by vacuum filtration, washed with a mixture of cold MeCN (30 mL) and DCM (30 mL), and dried to provide 15.1 g (69%) of **3** freebase Form III.

X-ray powder diffraction (XRPD) analysis for **3** was conducted on a diffractometer (PANalytical XPERT-PRO, PANalytical B. V., Almelo, Netherlands) using copper radiation (Cu K$\alpha$, $\lambda$ = 1.541874 Å). Samples were spread evenly on a zero-background sample plate. The generator was operated at a voltage of 45 kV and amperage of 40 mA. Slits were Soller 0.02 rad, antiscatter 1.0°, and divergence. Scans were performed from 2 to 40° 2θ with a 0.0167 step size. Data analysis was performed using X'Pert Data Viewer V1.2d (PANalytical B.V., Almelo, Netherlands).

Single crystals of **3** freebase Form III were obtained by vapor diffusion of heptane into saturated acetonitrile. **3** freebase Form III was weighed into a clear 4 mL vial, heated at 75°C with dropwise addition of acetonitrile until all the solids dissolved. The vial was then placed uncapped in a 20 mL vial containing heptane at about 20°C. Crystal growth was noted after 7 days; material was sent to Curia for single crystal XRD.

**Supporting Information (SI) Figures and Tables**

**SI Figure 1.** Characterization of **3** crystalline Form III

(a) X-Ray powder diffraction of **3** Form III

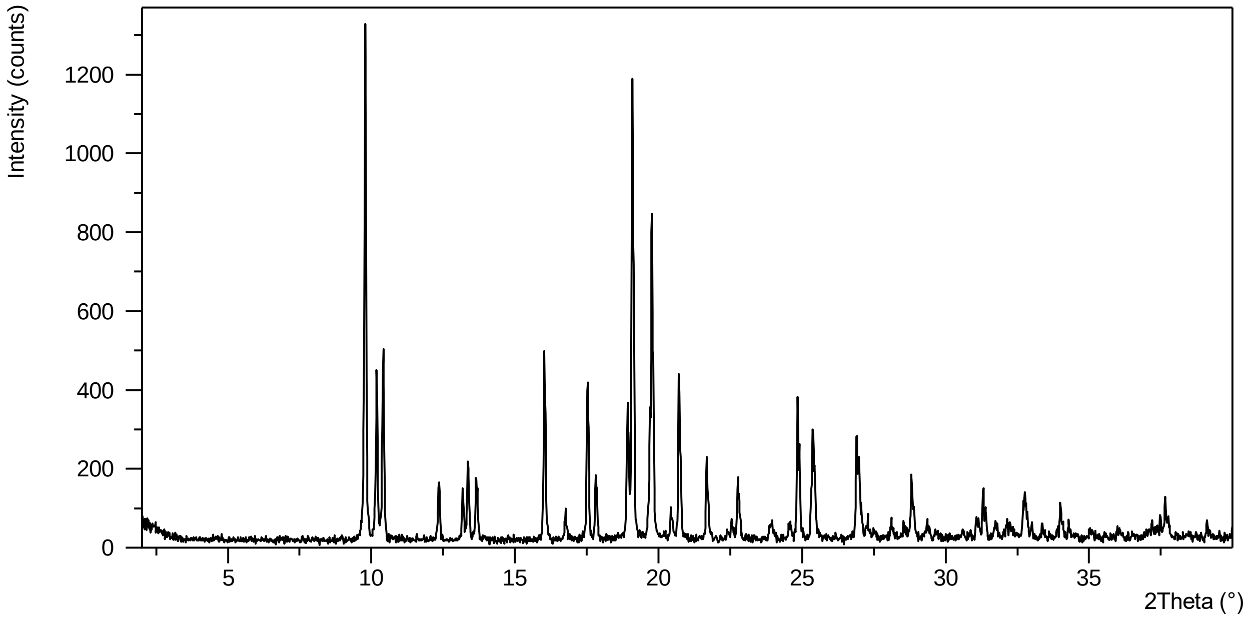


(b) X-ray structure of **3** Form III

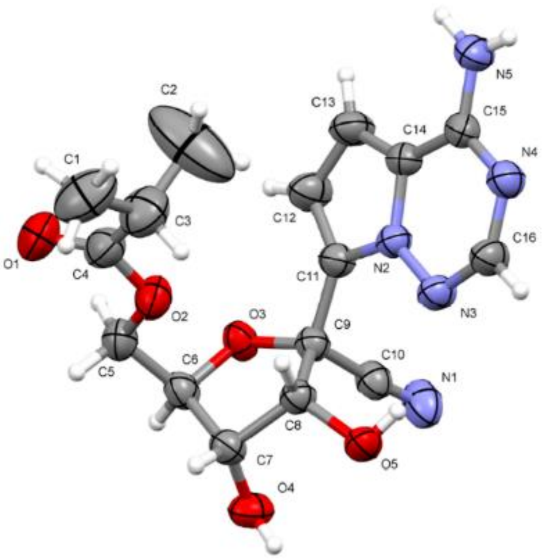

### Crystal data and data collection parameters

| Empirical formula | $C_{16}H_{19}N_5O_5$ |
|---|---|
| Formula weight (g mol$^{-1}$) | 361.36 |
| Temperature (K) | 299.64(10) |
| Wavelength (Å) | 1.54184 |
| Crystal system | orthorhombic |
| Space group | $P2_12_12_1$ |
| Unit cell parameters | |
| $a$ = 9.73530(10) Å | $\alpha$ = 90° |
| $b$ = 10.57480(10) Å | $\beta$ = 90° |
| $c$ = 17.3673(2) Å | $\gamma$ = 90° |
| Unit cell volume (Å$^3$) | 1787.94(3) |
| Cell formula units, $Z$ | 4 |
| Calculated density (g cm$^{-3}$) | 1.342 |
| Absorption coefficient (mm$^{-1}$) | 0.858 |
| F(000) | 760 |
| Crystal size (mm$^3$) | 0.27 × 0.13 × 0.13 |
| Reflections used for cell measurement | 6266 |
| $\theta$ range for cell measurement | 4.8780°–75.5770° |
| Total reflections collected | 8500 |
| Index ranges | $-11 \le h \le 11$; $-13 \le k \le 13$; $-19 \le l \le 21$ |
| $\theta$ range for data collection | $\theta_{min}$ = 4.896°, $\theta_{max}$ = 75.776° |
| Completeness to $\theta_{max}$ | 97.7% |
| Completeness to $\theta_{full}$ = 67.684° | 100% |
| Absorption correction | multi-scan |
| Transmission coefficient range | 0.976–1.000 |
| Refinement method | full matrix least-squares on $F^2$ |
| Independent reflections | 3590 [$R_{int}$ = 0.0161, $R_\sigma$ = 0.0189] |
| Reflections [ $I$>2σ($I$) ] | 3466 |
| Reflections / restraints / parameters | 3590 / 0 / 253 |
| Goodness-of-fit on $F^2$ | $S$ = 1.05 |
| Final residuals [ $I$>2σ($I$) ] | $R$ = 0.0357, $R_w$ = 0.0997 |
| Final residuals [ all reflections ] | $R$ = 0.0367, $R_w$ = 0.1007 |
| Largest diff. peak and hole (e Å$^{-3}$) | 0.274, −0.218 |
| Max/mean shift/standard uncertainty | 0.000 / 0.000 |
| Absolute structure determination | Flack parameter: 0.07(7)<br>Hooft parameter: 0.07(6)<br>Friedel coverage: 95.1% |

**SI Figure 2**. Intracellular metabolism of **3** (Panel A), **2** (Panel B), and **1** (Panel C) to the active metabolite **2-NTP** (solid line) compared to intracellular concentrations of parent nucleoside **2** (nuc, dotted line) in A549-hACE2 and NHBE cultures. Average **2-NTP** and **2** concentrations are indicated after dose-normalization to 1 µM.

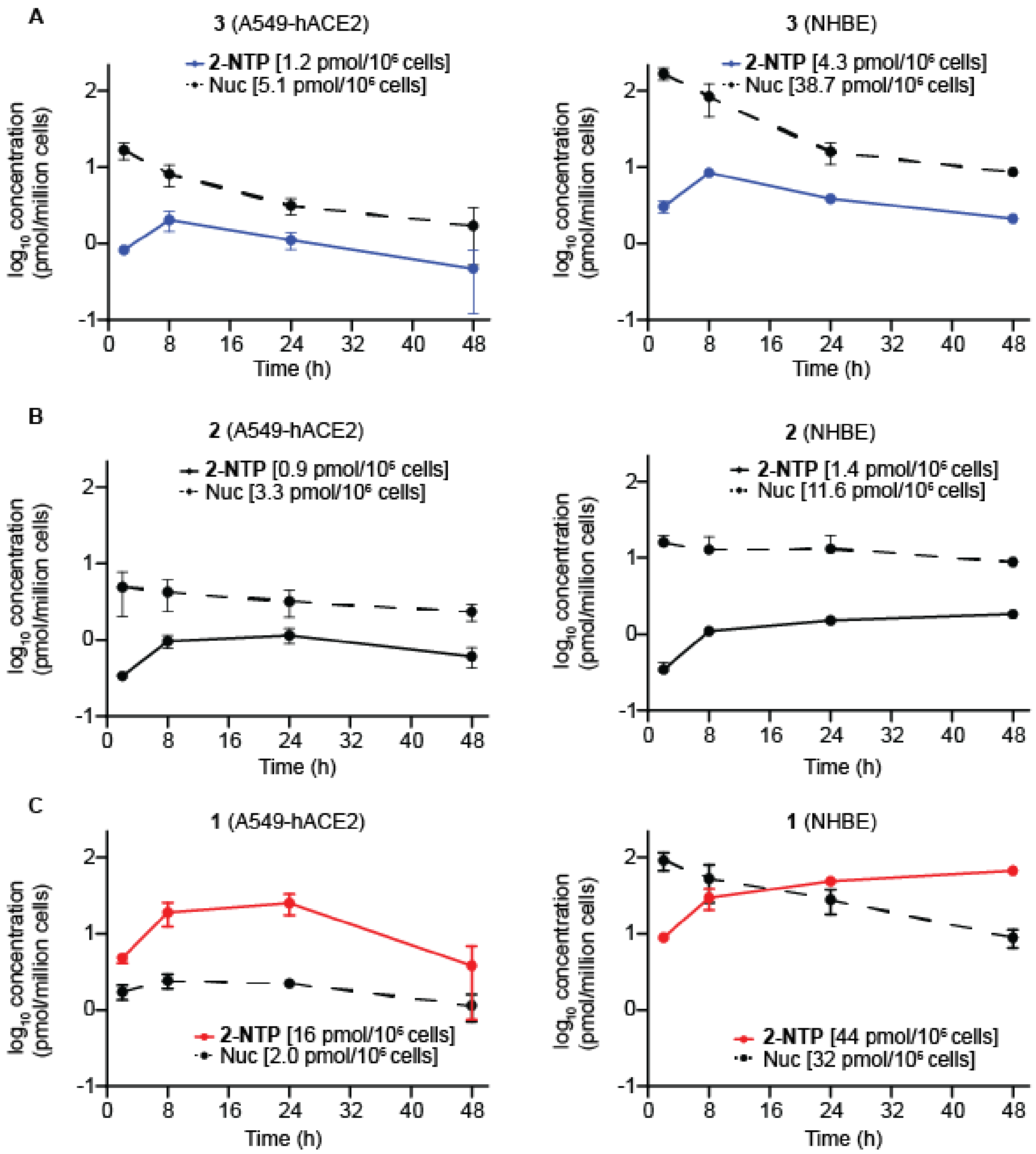

**SI Figure 3.** Extracellular concentrations of **3** (solid line) and parent nucleoside **2** (dotted line) following continuous incubation of A549-hACE2 (A) and NHBE (B) cultures with 10 µM of **3**.

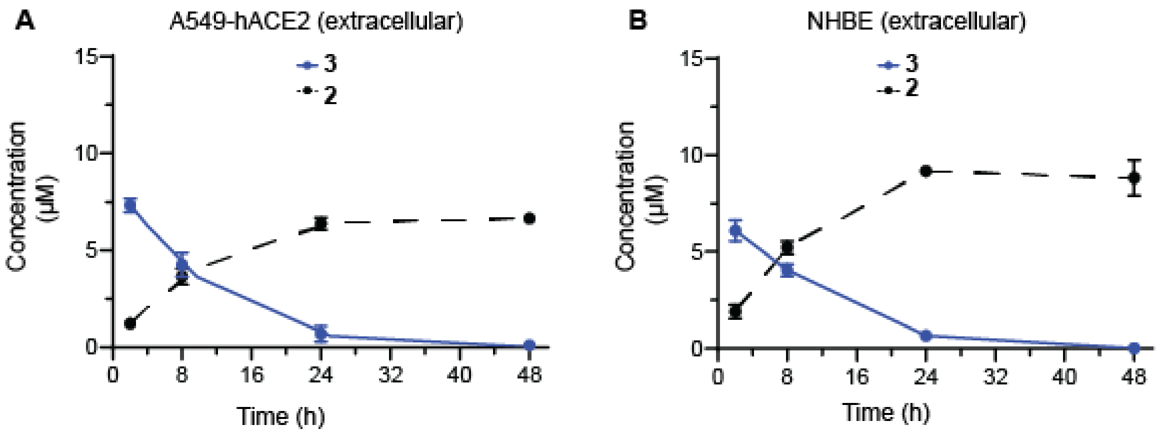

**SI Figure 4**. Absorbance spectra of test articles before and after treatment with recombinant human ADA1. Panel **A,** absorbance spectra of adenosine (positive control) and inosine (negative control) in the absence of ADA1 (solid lines) and after incubation with ADA1 (dotted lines) for 40 min. Panel **B**, absorbance spectra of **2**, **3** and **6** in the absence of ADA1 (solid lines) and after of incubation with ADA1 (dotted lines) for 58 min. Black solid line denotes absorbance spectra of 6 in the presence of ADA1 and ADA1 inhibitor EHNA.

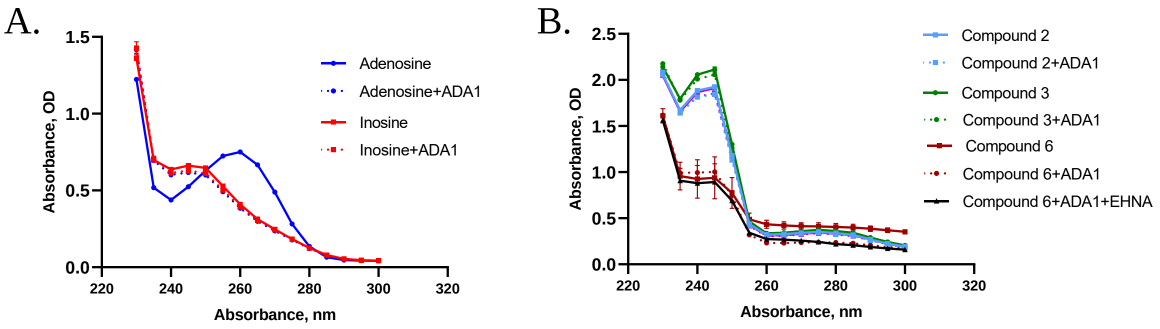

**SI Figure 5.** Nasal swab SARS-CoV-2 genomic RNA and Plaque Data. Antiviral effect of oral **3** on nasal swab samples in the African green monkey SARS-CoV-2 model. Panel **A**, Infectious virus in nasal swabs; Panel **B**, Genomic RNA in nasal swabs. LLOQ, lower limit of quantification. *p<0.05; **p<0.01; ***P<0.001; ****p<0.0001.

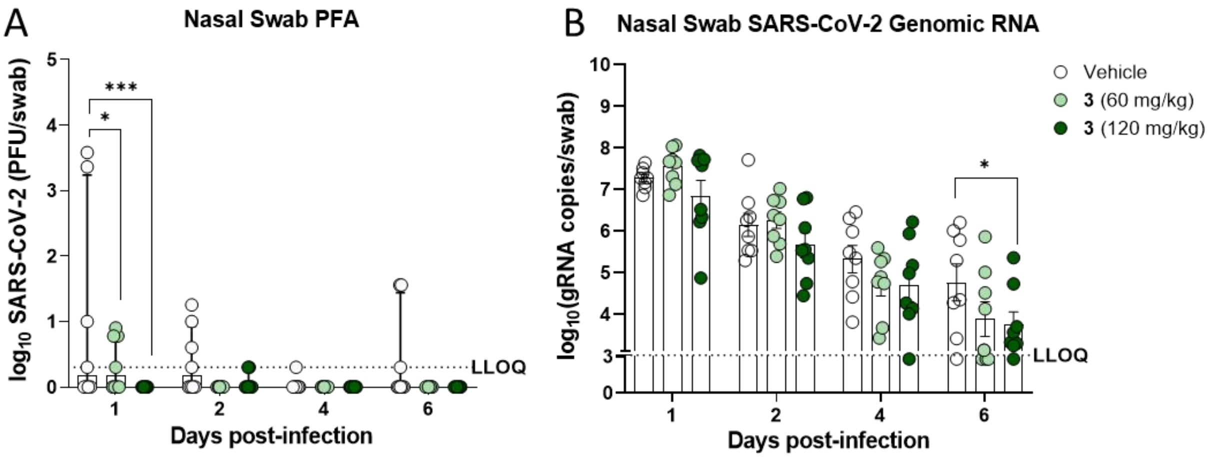

**SI Table 1.** Evaluation of **3** for potential mitochondrial toxicity.

| Mitochondria properties | Readouts | Compound $CC_{50}$ (µM)[a] | | |
|---|---|---|---|---|
| | | **3** | **2**[b] | Positive Control |
| Respiration<br>(3-day in PC-3) | Spare Respiratory Capacity<br>ATP level<br>Cell count | >100<br>>100<br>>100 | >100<br>>100<br>>100 | Chloramphenicol<br>4.5 ± 1.5<br>>50<br>>50 |
| Protein synthesis<br>(5-day in PC-3)[c] | COX-1 (mtDNA encoded)<br>SDHA (ncDNA encoded)<br>ATP level | <br>>100<br>>100<br>>100 | <br>>100<br>>100<br>>100 | Chloramphenicol<br>5.3 ± 0.4<br>>25<br>16 ± 2 |
| DNA synthesis<br>(10-day in HepG2) | Under different [compound concentration], amount of mtDNA (% of DMSO control) | <br>[0.4 µM], 100%<br>[4.0 µM], 100%<br>[40 µM], 100% | <br>[1.0 µM], 96 ± 32%<br>[10 µM], 90 ± 22%<br>[100 µM], 83 ± 12% | ddC<br>[0.2 µM], 97 ± 2%<br>[2.0 µM], 16 ± 5%<br>[20 µM], 0.64 ± 0.38% |

[a]All values represent the average ± standard deviation of at least three independent measurements. [b] Reference: Xu, 2021. [c] For mitochondrial protein synthesis analysis, two proteins cytochrome c oxidase subunit 1 (COX-1; encoded by mitochondrial DNA [mtDNA]) and succinate dehydrogenase (SDH-A; encoded by nuclear DNA [ncDNA]) are quantified simultaneously using immunocytochemistry.

**SI Table 2.** Adenosine Deaminase Inhibition and Substrate Potential

**(a)** Effect of test articles on catalytic activity of recombinant human ADA1

| **Test Article** | **$t_{1/2}$ adenosine[a] (min)** |
|---|---|
| DMSO, 1% | 4.7 ± 0.5 |
| **2** | 7 ± 2 |
| **3** | 7 ± 2 |
| **6** | 5 ± 1 |

[a]Half-life ($t_{1/2}$) of ADA1 catalyzed conversion of an adenosine to inosine in the absence (1% DMSO) and presence of a test article was monitored by change in the absorbance of adenosine measured at the wavelength of 265 nm. Values represent an average and a standard deviation of two independent determinations.

**(b)** Effect of recombinant human ADA1 on absorbance of test articles

| **Wavelength** | **Test Article** | **Absorbance of test article treated with ADA1[a]** | **% Absorbance change[b]** |
|---|---|---|---|
| 265 nm | Inosine | 0.312 ± 0.003 | -1.1 ± 0.7 |
| 265 nm | Adenosine | 0.367 ± 0.007 | 52 ± 1 |
| 245 nm | Inosine | 0.655 ± 0.008 | 0.2 ± 0.6 |
| 245 nm | **2** | 1.846 ± 0.006 | 7.0 ± 0.5 |
| 245 nm | **3** | 2.061 ± 0.006 | 6 ± 2 |
| 245 nm | **6** | 1.00 ± 0.07 | 3 ± 5 |
| 285 nm | Inosine | 0.0795 ± 0.0005 | -1 ± 0.6 |
| 285 nm | **2** | 0.308 ± 0.001 | 6.7 ± 0.4 |
| 285 nm | **3** | 0.333 ± 0.000 | 6 ± 2 |
| 285 nm | **6** | 0.21 ± 0.01 | 38 ± 3[c] |

[a]Absorbance of adenosine (positive control) and inosine (negative control) at 265 nm measured after 90 min of incubation with ADA1. Absorbance of test articles at 245 nm and 285 nm determined after 90 min incubation with ADA1 and compared to the absorbance of inosine at respective wavelengths. Values represent an average and a standard deviation of two independent determinations. [b]Percentage of absorbance change of adenosine (positive control), inosine (negative control) or a test article after 90 min incubation with ADA1, measured at respective wavelengths. Values represent an average and a standard deviation of two independent determinations. [c]Percentage of absorbance change independent on ADA1 treatment as similar change (38 ± 5%) observed in the presence of ADA1 inhibitor EHNA (Supplementary Figure 4B)

### SI Table 3

**(a)** SARS-CoV-2 viral parameters in bronchioalveolar lavage fluid following oral dosing of **3.**

| SARS-CoV-2 load in bronchioalveolar lavage fluid | | | | | | | |
|---|---|---|---|---|---|---|---|
| | | **Genomic RNA RT-qPCR** | | | **Infectious Virus** | | |
| **DPI** | **Dosing Group** | **mean log (copies/mL) (# BLQ)** | **mean diff.** | ***p* value** | **mean log (PFU/mL) (# BLQ)** | **mean diff.** | ***p* value** |
| 1 | Vehicle | 5.77 (0) | | | 4.33 (0) | | |
| | 60 mg/kg | 5.11 (0) | -0.67 | 0.168 | 3.05 (0) | -1.28 | 0.0452 |
| | 120 mg/kg | 4.55 (2) | -1.22 | 0.0039 | 1.78 (0) | -2.56 | <0.0001 |
| 2 | Vehicle | 6.42 (0) | | | 4.97 (0) | | |
| | 60 mg/kg | 4.41 (4) | -2.01 | <0.0001 | 3.32 (0) | -1.64 | 0.0076 |
| | 120 mg/kg | 4.20 (5) | -2.22 | <0.0001 | 1.19 (2) | -3.78 | <0.0001 |
| 4 | Vehicle | 5.22 (0) | | | 2.76 (0) | | |
| | 60 mg/kg | 3.82 (5) | -1.40 | 0.0009 | 0.52 (4) | -2.24 | 0.0002 |
| | 120 mg/kg | 3.63 (6) | -1.59 | 0.0002 | 0.0 (8) | -2.76 | <0.0001 |
| 6 | Vehicle | 4.84 (0) | | | 2.38 (1) | - | - |
| | 60 mg/kg | 3.43 (8) | -1.41 | 0.0009 | 0.04 (7) | -2.34 | 0.0001 |
| | 120 mg/kg | 3.43 (8) | -1.41 | 0.0009 | 0.0 (8) | -2.38 | <0.0001 |

BLQ, below limit of quantification; PFU, plaque forming units; DPI, day post infection.

**(b)** SARS-CoV-2 viral parameters in throat swabs following oral dosing of **3.**

| SARS-CoV-2 load in throat swab | | | | | | | |
|---|---|---|---|---|---|---|---|
| | | Genomic RNA RT-qPCR | | | Infectious Virus | | |
| **DPI** | **Dosing Group** | **mean log (copies/mL) (# BLQ)** | **mean diff.** | ***p* value** | **mean log (PFU/swab) (# BLQ)** | **mean diff.** | ***p* value** |
| 1 | Vehicle | 8.36 (0) | | | 4.41 (0) | | |
| | 60 mg/kg | 7.43 (0) | -0.93 | 0.0963 | 1.91 (1) | -2.50 | <0.0001 |
| | 120 mg/kg | 6.85 (0) | -1.51 | 0.0038 | 1.17 (0) | -3.24 | <0.0001 |
| 2 | Vehicle | 7.07 (0) | | | 1.21 (2) | | |
| | 60 mg/kg | 5.80 (0) | -1.27 | 0.0166 | 0.47 (4) | -0.74 | 0.0356 |
| | 120 mg/kg | 5.15 (0) | -1.92 | 0.0002 | 0.00 (8) | -1.21 | 0.0003 |
| 4 | Vehicle | 5.38 (0) | | | 0.59 (4) | | |
| | 60 mg/kg | 4.65 (0) | -0.73 | 0.2417 | 0.0 (8) | -0.59 | 0.1421 |
| | 120 mg/kg | 3.82 (0) | -1.56 | 0.0026 | 0.0 (8) | -0.59 | 0.1421 |
| 6 | Vehicle | 4.93 (1) | | | 0.86 (4) | | |
| | 60 mg/kg | 2.99 (3) | -1.94 | 0.0002 | 0.0 (8) | -0.86 | 0.0116 |
| | 120 mg/kg | 2.50 (5) | -2.43 | <0.0001 | 0.0 (8) | -0.86 | 0.0116 |

BLQ, below limit of quantification; PFU, plaque forming units; DPI, day post infection.

**(c)** SARS-CoV-2 viral parameters in nasal swabs following oral dosing of **3.**

| **SARS-CoV-2 load in nasal swab** | | | | | | | |
|---|---|---|---|---|---|---|---|
| | | **Genomic RNA RT-qPCR** | | | **Infectious Virus** | | |
| **DPI** | **Dosing Group** | **mean log (copies/mL) (# BLQ)** | **mean diff.** | ***p* value** | **mean log (PFU/swab) (# BLQ)** | **mean diff.** | ***p* value** |
| 1 | Vehicle | 7.27 (0) | | | 1.18 (4) | | |
| | 60 mg/kg | 7.55 (0) | 0.28 | >0.9999 | 0.42 (4) | -0.76 | 0.0262 |
| | 120 mg/kg | 6.84 (0) | -0.43 | 0.6814 | 0. 0 (8) | -1.18 | 0.0003 |
| 2 | Vehicle | 6.15 (0) | | | 0.55 (4) | | |
| | 60 mg/kg | 6.26 (0) | 0.11 | >0.9999 | 0.0 (8) | -0.55 | 0.1455 |
| | 120 mg/kg | 5.66 (0) | -0.49 | 0.5428 | 0.11 (6) | -0.44 | 0.3066 |
| 4 | Vehicle | 5.32 (0) | | | 0.08 (7) | | |
| | 60 mg/kg | 4.71 (0) | -0.61 | 0.3451 | 0.0 (8) | -0.08 | >0.9999 |
| | 120 mg/kg | 4.69 (1) | -0.63 | 0.3164 | 0.0 (8) | -0.08 | >0.9999 |
| 6 | Vehicle | 4.76 (1) | | | 0.54 (5) | | |
| | 60 mg/kg | 3.88 (3) | -0.88 | 0.0984 | 0.0 (8) | -0.54 | 0.1515 |
| | 120 mg/kg | 3.75 (1) | -1.01 | 0.0499 | 0.0 (8) | -0.54 | 0.1515 |

BLQ, below limit of quantification; PFU, plaque forming units; DPI, day post infection.

**(d)** SARS-CoV-2 viral parameters in respiratory tissues following oral dosing of **3.**

| **SARS-CoV-2 RNA load in respiratory tissues** | | | | |
|---|---|---|---|---|
| | | **Genomic RNA RT-qPCR** | | |
| **Tissue** | **Dosing Group** | **mean log (copies/g) (# BLQ)** | **mean diff.** | ***p* value** |
| Lower Bronchus | Vehicle | 6.11 (1) | | |
| | 60 mg/kg | 3.69 (8) | -2.42 | <0.0001 |
| | 120 mg/kg | 3.95 (7) | -2.16 | 0.0004 |
| Mainstem Bronchus | Vehicle | 6.71 (0) | | |
| | 60 mg/kg | 4.88 (4) | -1.83 | 0.0009 |
| | 120 mg/kg | 3.96 (6) | -2.75 | <0.0001 |
| Lower Lung | Vehicle | 6.25 (0) | | |
| | 60 mg/kg | 3.88 (7) | -2.37 | <0.0001 |
| | 120 mg/kg | 4.26 (6) | -1.99 | 0.0004 |
| Middle Lung | Vehicle | 6.92 (0) | | |
| | 60 mg/kg | 5.61 (1) | -1.31 | 0.0475 |
| | 120 mg/kg | 5.03 (2) | -1.89 | 0.0041 |
| Upper Lung | Vehicle | 6.04 (3) | | |
| | 60 mg/kg | 4.64 (4) | -1.40 | 0.1578 |
| | 120 mg/kg | 4.48 (5) | -1.56 | 0.1054 |
| Trachea | Vehicle | 5.58 (2) | | |
| | 60 mg/kg | 3.69 (8) | -1.89 | 0.0008 |
| | 120 mg/kg | 3.69 (8) | -1.89 | 0.0008 |

BLQ, below limit of quantification.

### Compound 3 and 5 Spectra

### HPLC : Compound 3

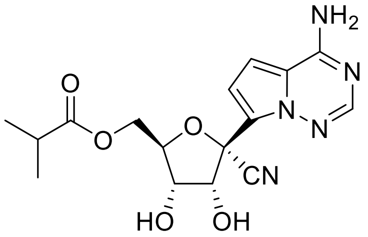


Method Info : Sample Bank Method - 2-98%B with 8.5 min gradient, A=Water + 0.1% TFA, B= Acetonitrile + 0.1% TFA; 1.5 mL/min; Column: Phenomenex Kinetex C18, 2.6u 100A, 4.6 x 100 mm; Instrument 1290II

Sample Info : Non-oral Nuc CoV pol inh; Rao Kalla

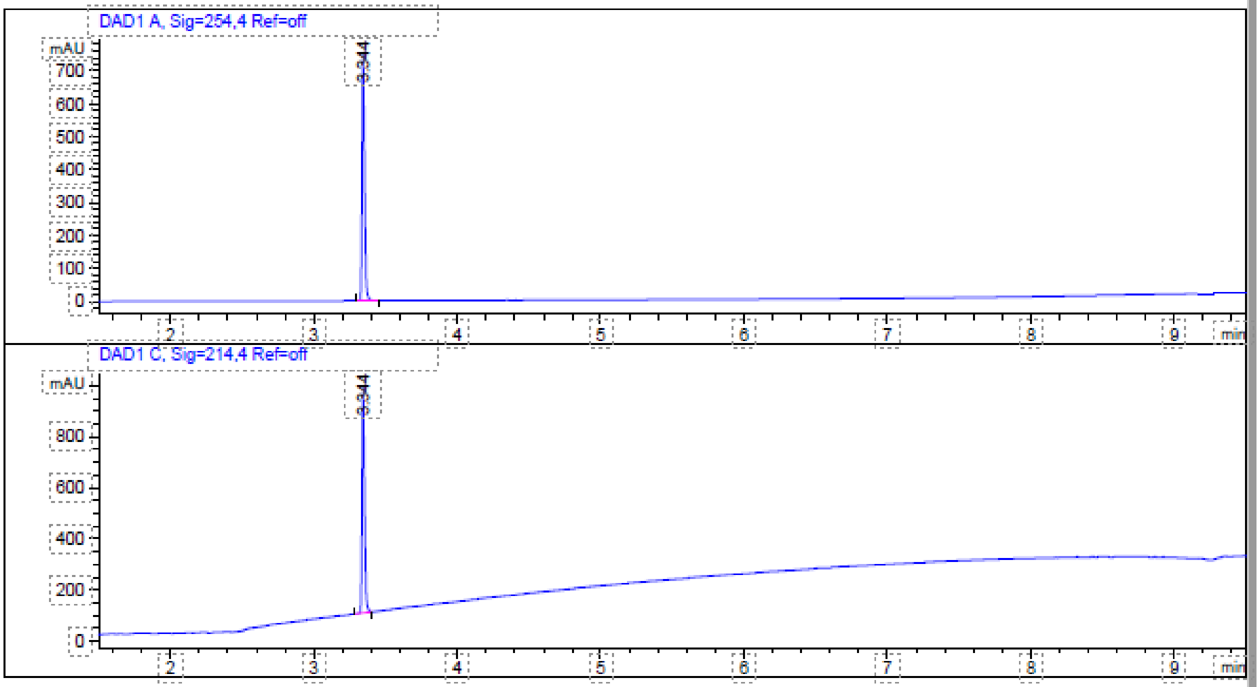


```
=====================================================================
                         Area Percent Report
=====================================================================

Sorted By             :      Signal
Multiplier            :      1.0000
Dilution              :      1.0000
Use Multiplier & Dilution Factor with ISTDs


Signal 1: DAD1 A, Sig=254,4 Ref=off
```

| Peak # | RetTime [min] | Type | Width [min] | Area [mAU*s] | Height [mAU] | Area % |
|---|---|---|---|---|---|---|
| 1 | 3.344 | BB | 0.0192 | 949.06555 | 754.39874 | 100.0000 |
| Totals : | | | | 949.06555 | 754.39874 | |

Signal 2: DAD1 C, Sig=214,4 Ref=off

| Peak # | RetTime [min] | Type | Width [min] | Area [mAU*s] | Height [mAU] | Area % |
|---|---|---|---|---|---|---|
| 1 | 3.344 | BB | 0.0190 | 1106.65112 | 888.36078 | 100.0000 |
| Totals : | | | | 1106.65112 | 888.36078 | |

==================================================================

*** End of Report ***

HPLC : Compound **5**

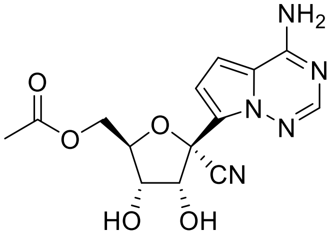


Method Info : Sample Bank Method - 2-98%B with 8.5 min gradient, A=Water + 0.1% TFA, B= Acetonitrile + 0.1% TFA; 1.5 mL/min; Column: Phenomenex Kinetex C18, 2.6u 100A, 4.6 x 100 mm; Instrument 1290II

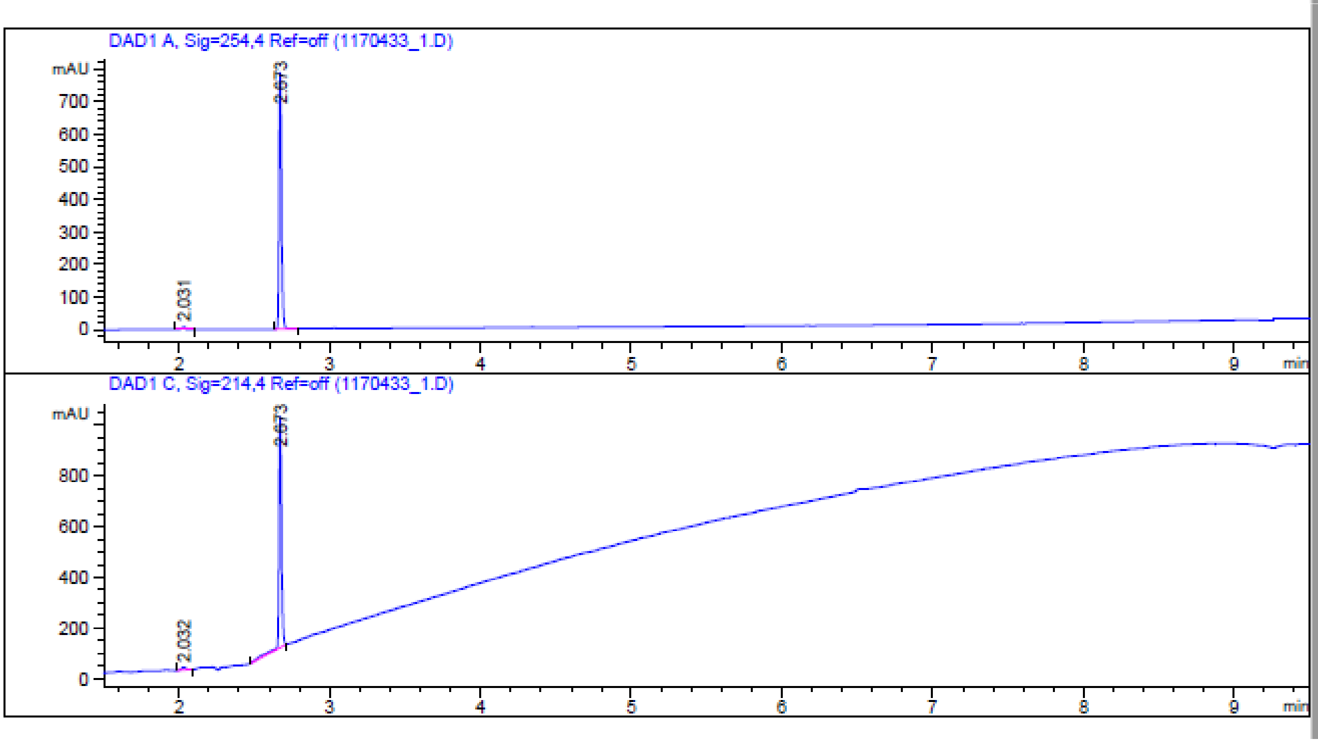

```
=====================================================================
                         Area Percent Report
=====================================================================

Sorted By             :      Signal
Multiplier            :      1.0000
Dilution              :      1.0000
Use Multiplier & Dilution Factor with ISTDs


Signal 1: DAD1 A, Sig=254,4 Ref=off


Peak RetTime Type  Width     Area       Height     Area
  #   [min]        [min]   [mAU*s]      [mAU]        %
----|-------|----|-------|----------|----------|--------|
   1   2.031 BB    0.0238   10.90270    6.78331   1.1912
   2   2.673 BB    0.0173  904.38593  790.38251  98.8088

Totals :                   915.28863  797.16582


Signal 2: DAD1 C, Sig=214,4 Ref=off

Peak RetTime Type  Width     Area       Height     Area
  #   [min]        [min]   [mAU*s]      [mAU]        %
----|-------|----|-------|----------|----------|--------|
   1   2.032 BB    0.0298   21.65358   10.20393   1.9516
   2   2.673 BB    0.0179 1087.85327  914.93555  98.0484

Totals :                  1109.50686  925.13948


=====================================================================
                          *** End of Report ***
```

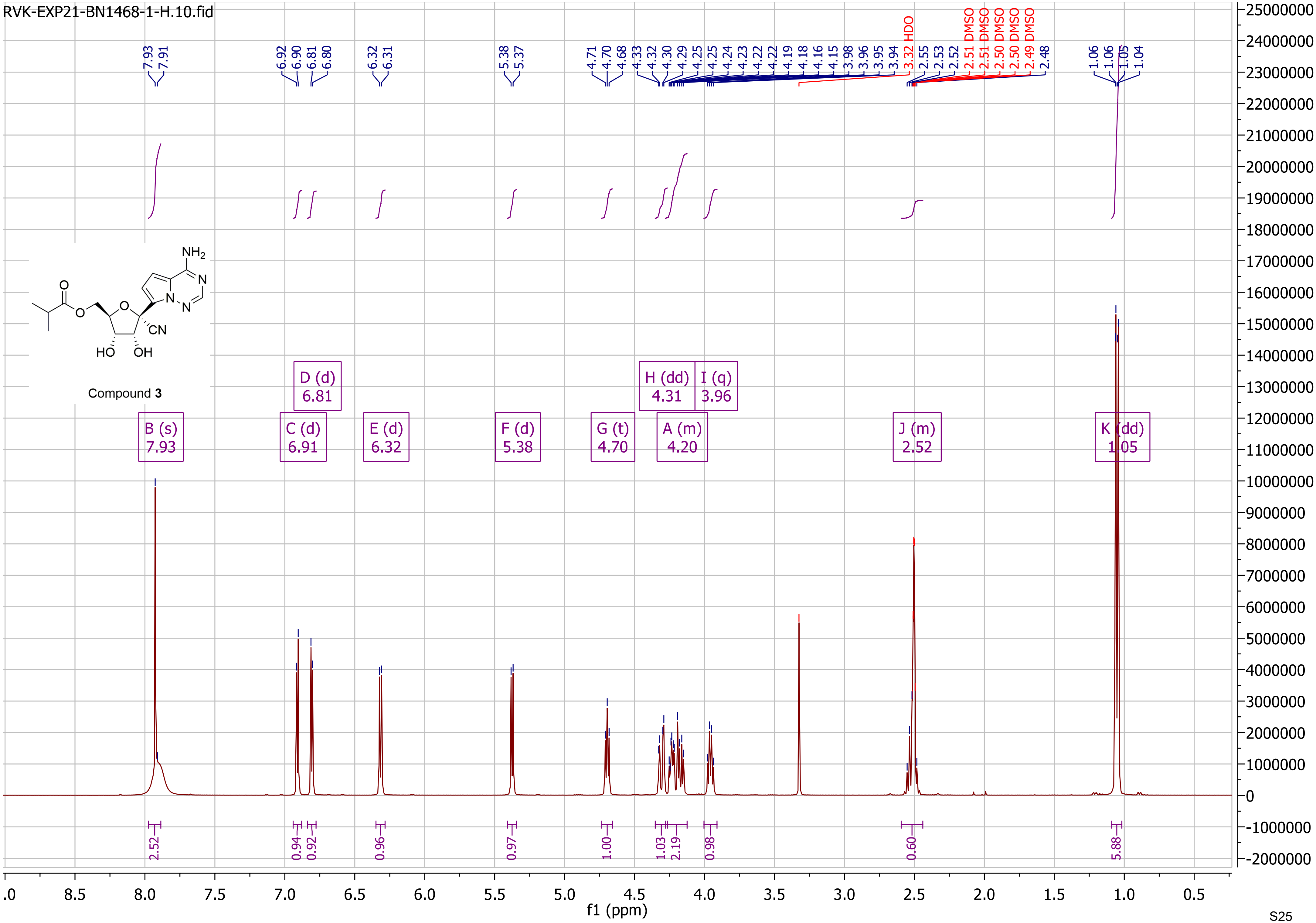

RVK-EXP21-BN1468-1-H.10.fid
Compound 3
B (s) 7.93
C (d) 6.91
D (d) 6.81
E (d) 6.32
F (d) 5.38
G (t) 4.70
H (dd) 4.31
A (m) 4.20
I (q) 3.96
J (m) 2.52
K (dd) 1.05
3.32 HDO
2.51 DMSO
2.50 DMSO
2.49 DMSO
f1 (ppm)

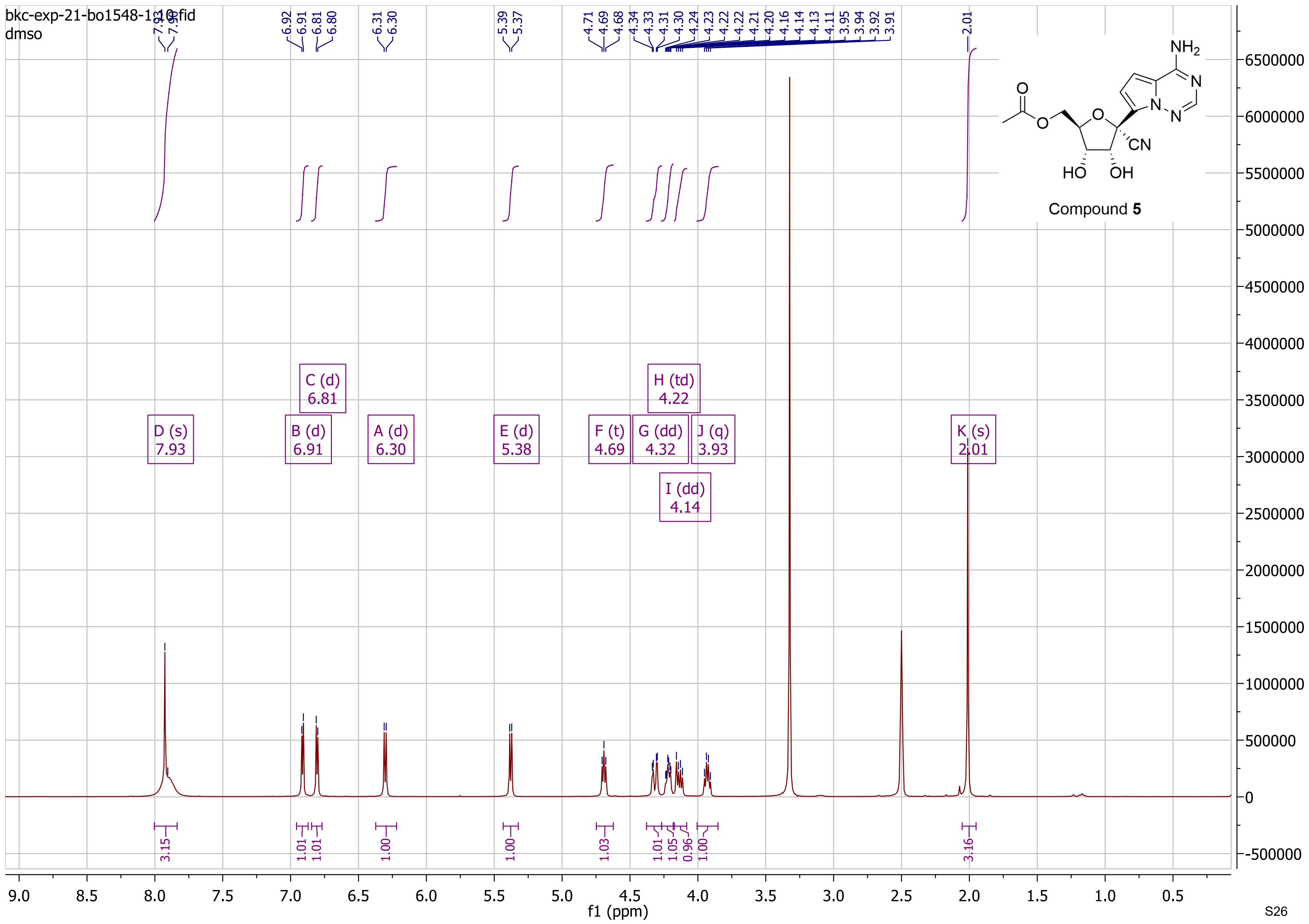

bkc-exp-21-bo1548-1-10.fid
dmso
Compound 5
D (s) 7.93
C (d) 6.81
B (d) 6.91
A (d) 6.30
E (d) 5.38
F (t) 4.69
G (dd) 4.32
H (td) 4.22
I (dd) 4.14
J (q) 3.93
K (s) 2.01
f1 (ppm)
